# Nucleus softens during herpesvirus infection

**DOI:** 10.1101/2025.01.03.631188

**Authors:** Aapo Tervonen, Visa Ruokolainen, Simon Leclerc, Katie Tieu, Sébastien Lyonnais, Henri Niskanen, Jian-Hua Chen, Alka Gupta, Minna U Kaikkonen, Carolyn A Larabell, Delphine Muriaux, Salla Mattola, Daniel E Conway, Teemu O Ihalainen, Vesa Aho, Maija Vihinen-Ranta

**Author notes:** Corresponding authors: (MVR), (VA). These authors contributed equally to this work (Aapo Tervonen, Visa Ruokolainen). These authors also contributed equally to this work (Vesa Aho, Maija Vihinen-Ranta).

## Abstract

Nuclear mechanics is remodeled not only by extracellular forces but also by internal modifications, such as those induced by viral infections. During herpes simplex virus type 1 infection, the nuclear structures undergo drastic reorganization, but little is known about how nuclear mechanobiology changes as a result. We show that the nucleus softens dramatically during the infection. To understand the phenomenon, we used advanced microscopy and computational modeling. We discovered that the enlarged viral replication compartment had a low biomolecular density, partially explaining the observed nuclear softening. The mobility of the nuclear lamina decreased, which suggests increased rigidity and an inability to induce softening. However, computational modeling supported by experimental data showed that reduced outward forces, such as cytoskeletal pull and intranuclear osmotic pressure acting both on and within the nucleus, can explain the decreased nuclear stiffness. Our findings reveal that during infection, the nucleus is subject to changes in multiple mechanical forces, leading to decreased nuclear stiffness.

**Author Summary:** DNA viruses take over the host cell nucleus, inducing dramatic structural modifications. There is currently very little knowledge of how the progression of viral infection modifies the mechanical properties of the nucleus, which are essential for various cellular processes, including gene expression and cell migration. Here, we show that the nucleus softens when herpesvirus infection progresses. We discovered that the viral replication compartment established in the central parts of the nucleus had a low biomolecular density, which may contribute to the nuclear softening. The shape and motion of the nuclear lamina suggested that it became more rigid, indicating that another mechanism was involved in the decreased elasticity. Our mechanical simulations and experiments showed that a reduction in outward forces, such as actin cytoskeleton pull or osmotic pressure, is the most likely factor in the nuclear softening. Our study provides new insights into the effects of DNA viruses on the mechanics of host cell nuclei, significantly expanding the knowledge of viral infection mechanobiology.

## Introduction

Mechanical forces have an essential role in many cellular functions. Cells sense and react to external forces, which can lead to various responses, such as changes in gene expression and cell differentiation [1,2]. Cells also generate intracellular forces, for example, to regulate cell shape and motion [3]. The forces are transmitted via cytoskeletal filaments and can be conveyed into the nucleus and to the chromatin via transmembrane proteins in the nuclear envelope (NE). In herpes simplex virus type 1 (HSV-1)-infected cells, expansion of the viral replication compartment (VRC) and virus-induced manipulation of nuclear structure and functions most likely lead to changes in the mechanical properties of the nucleus. However, little is known about the details of these changes.

The main mechanical components of the nucleus are the NE, with its underlying nuclear lamina, and the chromatin [4,5]. Structural changes in the organization of the NE, chromatin, or both often regulate nuclear stiffness [6,7]. The nuclear lamina, a 14-nm-thick filamentous network associated with the inner nuclear membrane [8], mainly comprises intermediate filament lamins divided into A/C and B types [9]. It reinforces the NE, connects to the cytoskeleton via the linker of the nucleoskeleton and cytoskeleton (LINC) complex, and interacts with the lamina-associated domains (LADs) of the chromatin [10–12]. The nuclear lamina, especially A-type lamins, has been shown to respond structurally to mechanical forces, and the amount of lamins is co-regulated by tissue rigidity [13,14]. Furthermore, reorganization of the lamina can change the ability of the nucleus to resist deformations and large strains [15]. The rigidity of chromatin, another regulator of nuclear stiffness, is modulated by changes in its compaction state [15–18] and by the chromatin crosslinker HP1α [19]. DNA damage and the LAD connections also contribute to the mechanical strength of the nuclei [7,20]. In addition, osmotic pressure in the nucleus has been suggested to affect its mechanical response [21]. Not only intranuclear components, but also cytoskeleton-mediated forces regulate nuclear mechanics. It has been shown that a force on the LINC complex protein nesprin-1 causes the nucleus to stiffen [22], and suppression of another LINC complex protein SUN2 results in nuclear softening [23].

Viral infections can cause drastic changes in nuclear function and structure. HSV-1 infection leads to the formation and expansion of spherical, membraneless VRCs for viral DNA replication and progeny capsid assembly [24–26], which is accompanied by reorganization of the host chromatin toward the nuclear periphery [27,28]. The VRC formation in HSV-1 and another herpesvirus, human cytomegalovirus, infection has properties similar to liquid-liquid phase separation. However, other compartmentalization mechanisms might also be included [29–31]. The substantial chromatin condensation toward the nuclear periphery regulates the diffusion of newly formed capsids through the chromatin to the NE [32–36], where the first step of nuclear egress, viral primary envelopment through the inner nuclear membrane, is assisted by the viral nuclear egress complex and the disintegration of the nuclear lamina [37–39].

In this study, we analyze HSV-1 infection-induced changes in the nuclear biomechanics of host cells by using various experimental methods, including atomic force microscopy (AFM), fluorescence lifetime imaging microscopy (FLIM), volume electron microscopy (EM), and cryo-soft X-ray tomography (SXT), as well as by using computational modeling. Our data show that the nuclei become softer as the infection progresses, while the nuclear structure undergoes profound changes and the nuclear lamina is modified. The modeling and experimental data suggest that, in addition to these changes, cytoskeletal pull on the nucleus and osmotic pressure also contribute to the softening.

## Results

### Nuclear stiffness decreases as the infection proceeds

To analyze how HSV-1 infection alters nuclear stiffness, we measured the mechanical response of Vero cell nuclei to compressive force using AFM [40]. The nuclei were subjected to indentation by a spherical tip of the AFM cantilever. The measurement of the bending of the cantilever with a known spring constant provides the force applied to the nucleus (Figs 1A and 1B, see also S1 Movie), which allows quantification of its elastic modulus, i.e., nuclear stiffness [41,42]. The measurements were conducted in intact cells, so although almost all the volume under the tip was occupied by the nucleus, thin layers of cytoplasm remained above and below the nucleus.

**Fig 1.**
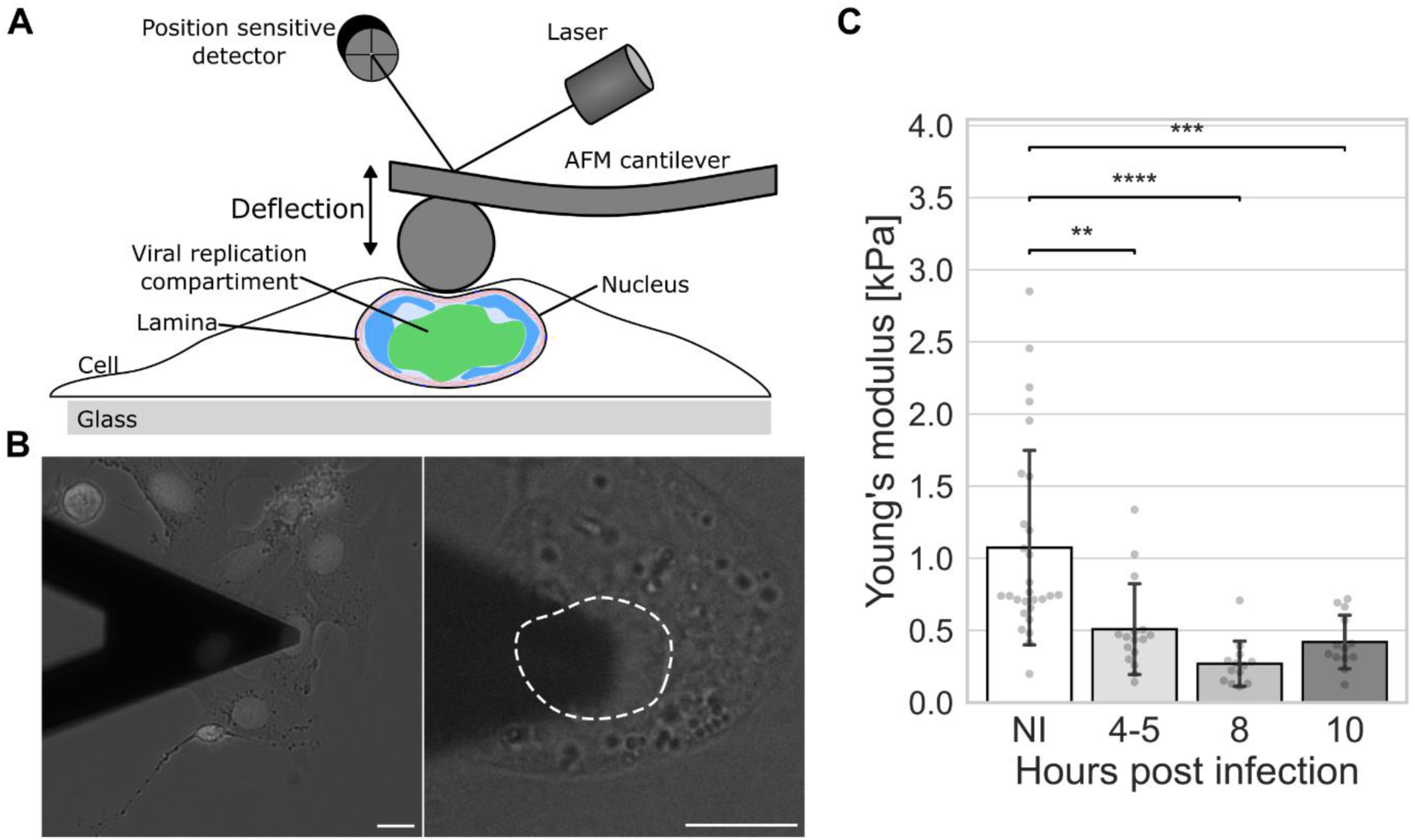
Nuclear stiffness decreases in HSV-1 infection. **(A)** Schematic side view of an atomic force microscope (AFM) cantilever with a colloidal probe over the nuclear region of a cell. **(B)** Brightfield microscopy images of an infected cell underneath the AFM cantilever at 8 hpi. The nuclear periphery is demarcated by a dotted white line. Infected Vero cells were identified by the presence of ICP4 by epifluorescence. See also S1 Movie. Scale bar, 10 μm. **(C)** AFM analysis of nuclear stiffness performed on noninfected and HSV-1 ICP4-EYFP-infected cells at 4-5, 8, and 10 hpi (n=28, 16, 13, and 13, respectively). The error bars show the standard deviation. Statistical significance was determined using Tukey’s test, and the significance values are denoted as **** (p < 0.0001), *** (p<0.001), or ** (p < 0.01). Nonsignificant differences (p ≥ 0.05) are not labeled.

There was a large variation in the stiffness of noninfected cell nuclei, with values ranging from 200 to 3,000 Pa. Compared to noninfected cells, the mean stiffness during the infection was lower at 4 hours post infection (hpi) and kept decreasing until 8 hpi, when Young’s modulus was only about one fourth of the noninfected cell value (Fig 1C). Finally, there was a minor recovery at 10 hpi.

Our findings show that the progression of HSV-1 infection softens the nucleus. To understand this phenomenon, we next studied how the infection modifies the mechanical components of the nucleus, such as chromatin and the nuclear lamina.

### The molecular density of the nuclear center decreases in the infection

In the lytic HSV-1 infection, the nuclear import of viral genomes is followed by the emergence of distinct punctate VRCs at early infection (1**–**5 hpi) [43]. Later in infection (8**–**12 hpi), the growth and fusion of the VRC foci result in an enlarged VRC, marginalization of host cell chromatin adjacent to the nuclear envelope, and expansion of the nuclear volume [27,28,44].

SXT imaging and analysis of infected mouse embryonic fibroblast (MEF) cells showed that biomolecular density, quantified by the linear absorption coefficient (LAC), in the central areas of the infected cell nuclei occupied by the VRC (Figs 2A and 2B, see also S2 Movie) was lower than that in the euchromatin areas of the noninfected cells (Fig 2C). The mean LAC values were 0.30, 0.25, and 0.24 μm^-1^ for noninfected cells and infected cells at 4 and 8 hpi, respectively. Notably, the density of the perinuclear chromatin in infected cells remained similar to the density of the heterochromatin in noninfected cells, with the mean LAC value of 0.33 μm^-1^ for every infection time point (Fig 2D).

**Fig 2.**
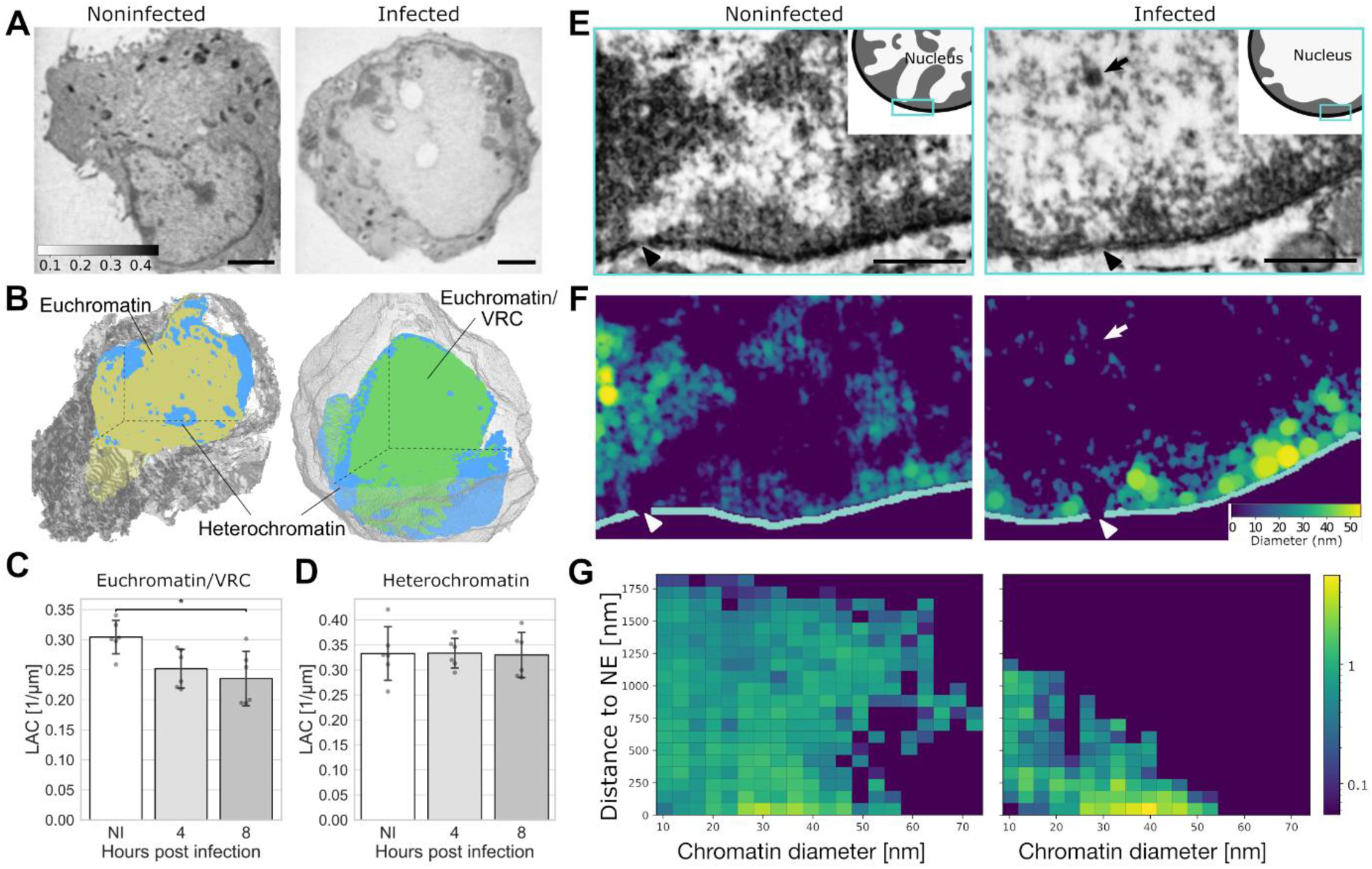
The enlarged viral replication compartment has a low biomolecular density. **(A)** Cryo-soft X-ray (SXT) tomography ortho-slices of noninfected (NI) and infected mouse embryonic fibroblast (MEF) cells at 8 hpi. Molecular densities are presented in a continuous grey scale representing linear absorption coefficient values ranging from 0.1 to 0.4 μm^-1^ (calibration bar). Scale bars, 2 μm. **(B)** Cross-sections of 3D SXT reconstructions of a noninfected and an infected cell. The high-density region (heterochromatin) is shown in blue, the low-density region (euchromatin) in yellow, the viral replication compartment (VRC/euchromatin) area in green, and the cytoplasm in grey. See also S2 Movie. Quantitative analysis of linear absorption coefficient (LAC) of the **(C)** low-density and **(D)** high-density region in noninfected and infected cells at 4 and 8 hpi (n=6). **(E)** Representative cross-sections of focused ion beam scanning electron microscopy (FIB-SEM) images of noninfected and infected MEF cells at 8 hpi. The viral capsid (arrows) and nuclear pore complexes (arrowheads) are shown. Scale bars, 0.5 µm. See also S3 Movie. **(F)** Reconstruction of chromatin fiber thickness and **(G)** chromatin density as a function of the distance from the nuclear envelope in noninfected and infected cells. The color scale in **(F)** indicates high and low chromatin thickness (yellow-blue), and in **(G)** high and low density (yellow-blue). The error bars show the standard deviation. Statistical significance was determined using Tukey’s test, and the significance values are denoted as * (p<0.05). Nonsignificant differences (p ≥ 0.05) are not labeled.

To further study the three-dimensional distribution and organization of chromatin in the nucleoplasm of MEF cells, we used focused ion beam scanning electron microscopy (FIB-SEM) and the ChromEM chromatin labeling method [45]. This approach confirmed the presence of dense chromatin near the NE in infected cells at 8 hpi (Fig 2E). Notably, finger-like protrusions of dense chromatin towards the center of the nucleus were less pronounced in infected cells. However, small and dense chromatin aggregates were still detected in the central area of the nucleus where the VRC and viral capsids were located (Figs 2F and 2G, see also S3 Movie). This can be cellular chromatin remaining in the VRC region or replicated viral genomes [46]. The ChromEM labeling can detect viral DNA, as shown by the intense labeling of DNA-containing capsids (S1 Fig).

In summary, the enlarged low-density region emerging during HSV-1 infection had a lower biomolecular density than euchromatin in the central parts of noninfected cell nuclei. The density decreased as the infection proceeded, which might have implications for the nuclear softness during the infection. The marginalized chromatin of the infected cell was not denser than the control cell heterochromatin.

### The nuclear lamina motion decreases and the nuclear membrane tension increases in the infection

Since the nuclear lamina is an essential mechanical structure for the nucleus, we used lamin A/C chromobody to assess differences in the nuclear lamina shape and motion in noninfected and infected living Vero cells [47]. We hypothesized that the mechanical stiffness of the nuclear lamina is linked to its shape and mobility, and that an increase (or decrease) in internal pressure would cause the nucleus to bulge (or wrinkle).

To analyze the lamina shape, it was segmented, skeletonized, and discretized. Local regions of positive and negative curvature with respect to the nuclear interior were then quantified (Fig 3A). The proportion of negative curvature regions was smallest at 8 hpi, but the change was not statistically significant (Fig 3B). Next, the extent of lamina fluctuations was analyzed by taking the maximum intensity projection of the segmented and skeletonized lamina over the time series and calculating its area (Fig 3C). The motion of the nuclear lamina, quantified as the area covered by the lamina skeleton during the time series and normalized by the length of the lamina, was significantly smaller at 8 hpi than in noninfected cells (Fig 3D). This shows that the motion of the lamina decreases in infection, indicating higher tension or reduced mechanical dynamics. Finally, nuclear membrane tension was measured by the ER Flipper-TR probe and FLIM [48]. Changes in the fluorescence lifetime indicated increased tension in the NE at 12 hpi compared to noninfected cells, although at 8 hpi the membrane tension was similar to that in the noninfected cells (Figs 3E and 3F).

**Fig 3.**
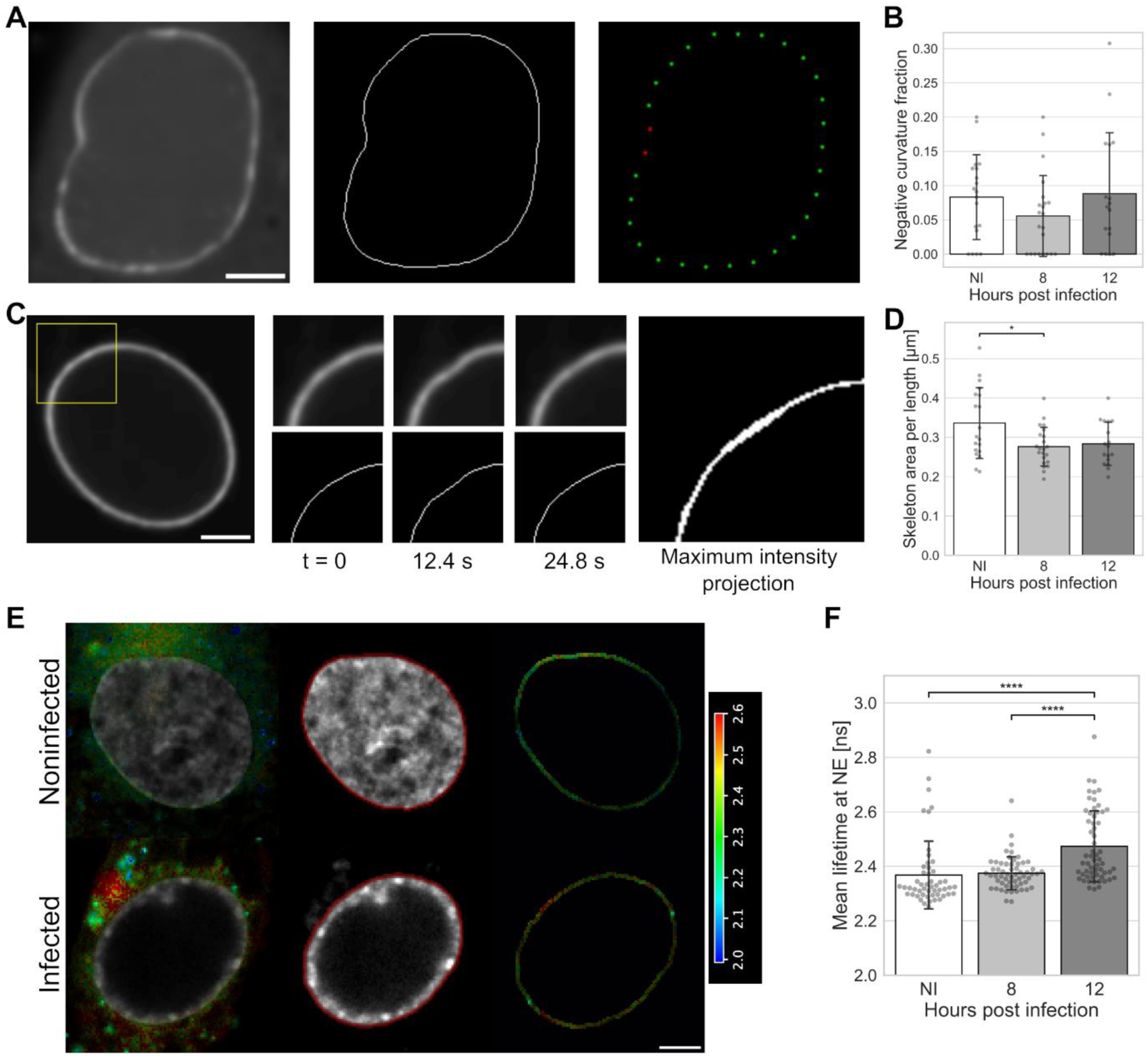
The motion of the nuclear lamina decreases in infection. **(A)** Nuclear lamina labeled by lamin A chromobody, skeletonization of the lamina, and discretized lamina points showing regions of positive (green) and negative curvature (red) in infected Vero cells at 8 hpi. **(B)** The fraction of negative curvature points for noninfected cells and cells imaged 8 and 12 hpi (n=19, 22, and 17, respectively). **(C)** A zoom-in of the nuclear lamina, showing its motion and maximum intensity projection of the skeleton during 24.8 s. Scale bars, 5 µm. **(D)** Nuclear lamina mobility, measured as the area covered by the skeleton during one minute for noninfected cells and cells imaged 8 and 12 hpi (n=19, 22, and 17, respectively). **(E)** Nuclear envelope (NE) tension analysis by fluorescence lifetime imaging microscopy (FLIM) of noninfected and HSV-1 17+ wild-type virus-infected Vero cells at 12 hpi using fluorescence lifetime-reporting membrane tension probe ER Flipper-TR. A fast FLIM image presented in a pseudocolor scale representing average photon arrival times in each pixel, ranging from 2.0 to 2.6 ns (calibration bar). Representative cells with average lifetime distribution, segmented NE (red) around the Hoechst signal (grey), and average lifetime at the NE region are shown. Scale bar, 10 µm. **(F)** The longer lifetime component of the ER Flipper probe, responsive to the membrane tension, in noninfected and infected cells at 8 and 12 hpi (n=51, 58, and 59, respectively). The error bars show the standard deviation. Statistical significance was determined using Tukey’s test, and the significance values are denoted as * (p<0.05) or **** (p<0.0001). Nonsignificant differences (p ≥ 0.05) are not labeled.

Finally, we analyzed the significance of lamins in the infection-induced nuclear growth in Vero cells by using overexpression of A-type lamins and the presence of truncated and farnesylated lamin A protein, progerin [49]. Confocal microscopy showed that the progression of infection and overexpression of lamin A/C increased nuclear volume, whereas the presence of progerin reduced it (S2 Fig).

### SUN2 expression decreases in infection

The nuclear lamina is one of the significant determinants of the mechanical behavior of the nucleus. Thus, to explore the transcriptional activity of lamina-associated proteins during infection, we performed global run-on sequencing (GRO-Seq) in Vero cells. The gene ontology analysis (GO) of GRO-Seq data according to the PANTHER classification system for the GO biological process (http://www.pantherdb.org) identified three main categories: protein localization to the NE (GO:0090435), nuclear migration (GO:0007097), and NE organization (GO:0006998) (Figs 4A and 4B, see also S1 Table).

**Fig 4.**
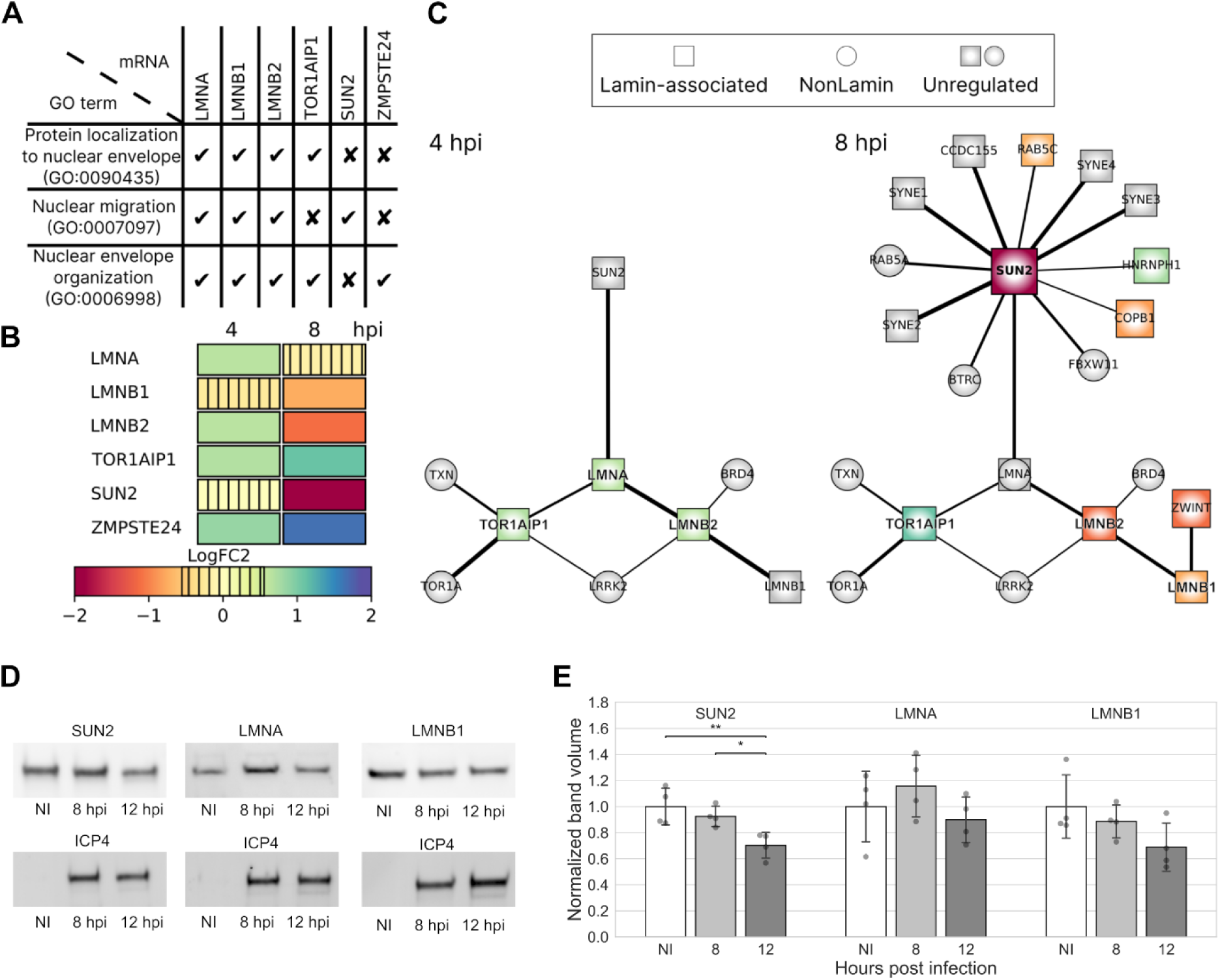
The amount of SUN2 decreases in infection. Global run-on sequencing (GRO-seq) analysis of nascent RNA levels of nuclear lamins and lamin-associated proteins in infected Vero cells at 4 and 8 hpi. **(A)** GO term classification for lamin-associated processes and **(B)** regulation of proteins. The color scale indicates upregulation (green-blue), downregulation (orange-red), or low change of gene transcription (vertical stripes) compared to noninfected control cells. **(C)** The genes encoding for lamins and lamin-associated proteins (square nodes) and cellular interactor proteins not associated with lamins (circular nodes) are shown. The upregulated (green-blue) or downregulated (red-orange) transcripts in response to the infection are visible together with unregulated interacting transcripts (grey). The node size correlates with the logarithmic fold change (logFC) of regulation, and the thickness of black lines between the nodes is proportional to the interaction in the STRING database (https://string-db.org/). **(D**) Representative western blots showing the expression of SUN2, lamin A, and lamin B in noninfected and infected cells at 8 and 12 hpi. Viral ICP4 was used as an infection marker. **(E)** Quantitative analysis of western blots (n = 4). The error bars show the standard deviation. Statistical significance was determined using Tukey’s test, and the significance values are denoted as ** (p<0.01) or * (p<0.05). Nonsignificant differences (p ≥ 0.05) are not labeled.

At 4 hpi, the genes encoding lamins A/C and B2 were upregulated compared to noninfected control cells, whereas at 8 hpi, the genes encoding lamins B1 and B2 were downregulated. Notably, transcription of SUN2, one of the essential proteins of the LINC complex and the key regulator of nucleus-cytoskeleton connections [50], was strongly downregulated at 8 hpi. The zinc metallopeptidase STE24 (ZMPSTE24), a protein needed during the preprocessing of lamin A [51], was upregulated at 4 and 8 hpi. This protein has a potent antiviral activity, inhibiting the replication of many enveloped DNA and RNA viruses [52,53]. At 4 and 8 hpi, torsin-1A-interacting protein 1 (TOR1AIP1, also known as LAP1), a protein involved in the connection between the nuclear membrane and the nuclear lamina [54,55], was upregulated. The downregulation of SUN2 at 8 hpi could decrease overall interactions between SUN2 and LINC complex components, including nesprin family proteins encoded by SYNE 1–4 genes [56]. The clustering of genes (https://string-db.org/) (Fig 4C and S1 Table) confirmed an interaction between SUN2 and lamin A/C, which further interacted with lamin B2. In addition, lamin A/C interacted with the upregulated TOR1AIP1.

Consistent with the GRO-seq data, western blot analysis showed that HSV-1 infection downregulated SUN2 expression at 8 hpi and, especially, at 12 hpi compared to noninfected cells. Our results are consistent with previous studies indicating a decrease in SUN2 protein levels at late stages of HSV-1 infection [57]. Protein expression levels of lamin A at 8 and 12 hpi were not significantly different from those of noninfected cells. The infection led to a slight downregulation of lamin B2 expression at 8 and 12 hpi, although this was not statistically significant (Figs 4D and 4E, see also S3 Fig). Altogether, our findings show that progression of infection led to downregulation of the key regulator of cytoskeleton–nucleus force transmission, the LINC complex protein SUN2, but did not significantly alter the expression of nuclear lamina proteins.

### Computational modeling of nuclear mechanics

Our experimental data indicated that the nuclear lamina becomes less mobile, which suggests that the NE becomes stiffer during the infection. However, AFM indentation measurements suggested that nuclear stiffness decreases during the infection. To understand this seeming contradiction, we built a computational model of the nucleus to simulate scenarios that could lead to the observed changes. The main model assumptions are: (1) the mechanical properties of the nucleus are governed by the nuclear envelope, including nuclear lamina, chromatin, and outward forces, representing actin cytoskeleton pull on the nucleus and osmotic pressure difference; (2) viscous changes in the nuclear structures occur on a time scale too slow to affect the fast AFM measurement; and (3) the VRC is soft and does not interfere with the AFM measurement.

The model consisted of two components that determine the main mechanical properties of the nucleus: the NE, which is considered to include the nuclear lamina, and chromatin. The NE was modeled as a thin 3D triangulated shell, and 46 separate polymers inside the shell represented the chromatin. Furthermore, connections between the two, the LADs, and transient cross-links within the chromatin were included in the model. Additionally, an outward force was included to represent the combined effects of outward cytoskeletal forces on the NE and osmotic pressure. The NE in the model includes all the mechanics related to changes in nuclear surface area, while the chromatin and outward forces together describe the mechanics associated with nuclear volume. Initial simulations indicated that both the nuclear area- and volume-related components contribute to the total nuclear stiffness. Finally, to model the VRC, another shell, similar to the NE, was added inside the nucleus and grown to an appropriate size to marginalize the chromatin towards the periphery (Fig 5A and S4 Movie). Model parameters for the noninfected nucleus were taken from the literature or fitted by replicating the AFM experiments (S5 Movie). The developed model has certain limitations. Combining the cytoskeletal pull and osmotic pressure into the outward forces reduces the ability of the model to predict the specific cause of the decreased outward force. Additionally, the model does not distinguish between heterochromatin and euchromatin, so it cannot describe the mechanical differences between them. However, adding these features would increase the number of unknown parameters that need to be estimated, thereby reducing the usability of the model.

**Fig 5.**
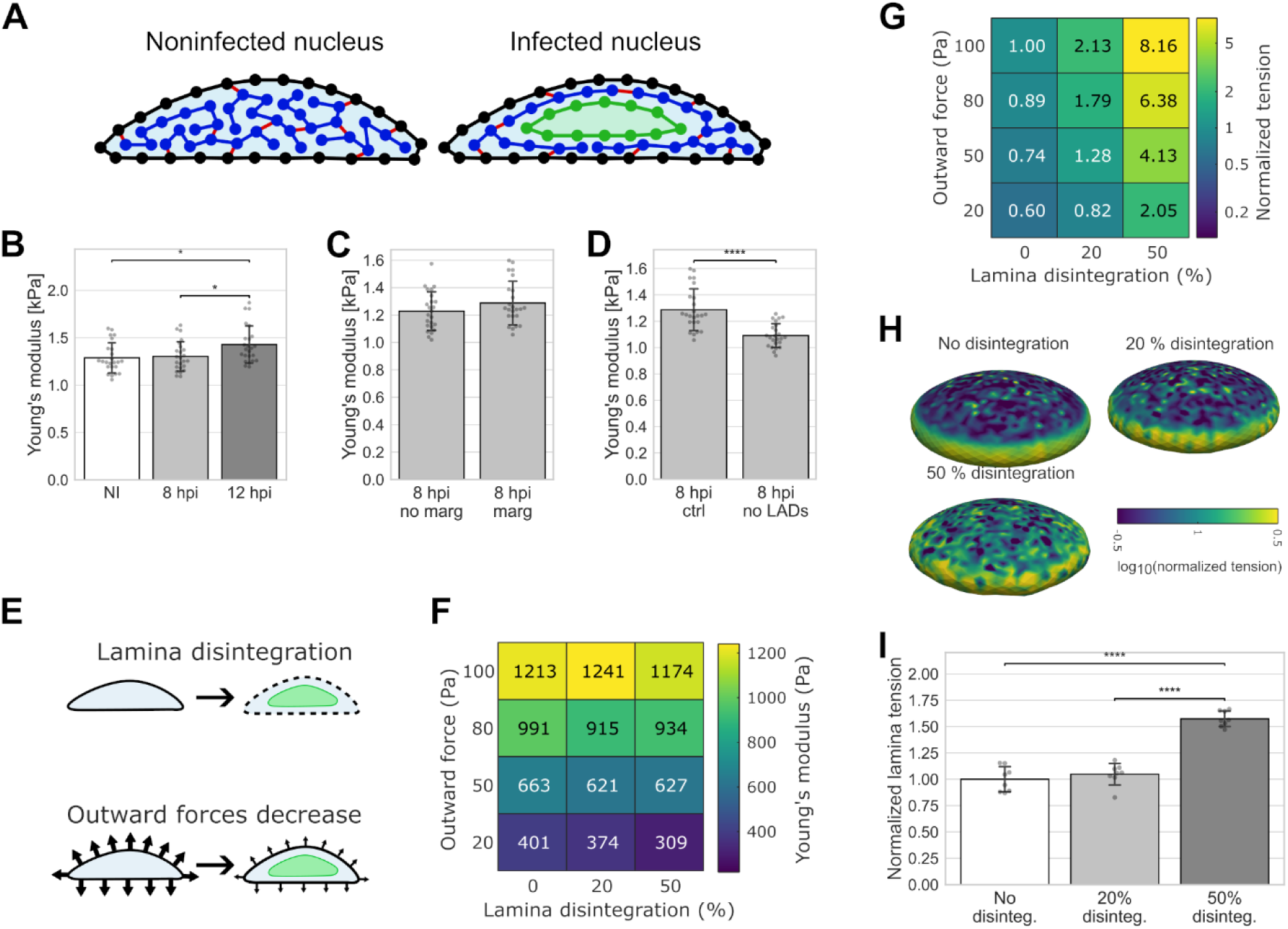
Changes in mechanical parameters alter the nuclear stiffness. **(A)** The setup for the computational model shows a noninfected and an infected cell nucleus. The nuclear envelope (black line), chromatin polymers (blue), envelope-chromatin and chromatin-chromatin connections (red lines), and the viral replication compartment (green line) are shown. **(B)** Simulated Young’s moduli for the noninfected (NI) and infected cells (8 and 12 hpi), **(C)** for the infected nuclei (8 hpi) without and with chromatin marginalization close to the nuclear envelope, and **(D)** for the infected nuclei with and without lamina-associated domains (LADs) (n=24). **(E)** Simulation of two changes in the nuclear mechanical parameters: lamina disintegration and decrease in the outward forces. **(F)** Simulated Young’s moduli as a function of parameters in infected cell nuclei at 8 hpi (n=8). Significances are shown in S4 Fig. **(G)** Normalized membrane tension as a function of lamina disintegration and outward forces (n=8). Significances are shown in S4 Fig. **(H)** Representative examples of lamina forces with no lamina disintegration and disintegration of 20% and 50%. **(I)** The sum of normalized lamina forces on the top surface of the nuclei without or with 20% or 50% disintegration (normalized to the mean value with no disintegration) (n=8). The error bars show the standard deviation. Statistical significances were determined using Tukey’s test (b, i) or Student’s t-test (c, d), and the significance values are denoted as **** (p<0.0001) or * (p<0.05). Nonsignificant differences (p ≥ 0.05) are not labeled.

To study nuclear mechanics during HSV-1 infection, various model parameters were iteratively adjusted to observe their impact on simulated AFM results. Based on the observed low molecular density (Figs 1A-1C), we hypothesized that the VRC is very soft and therefore disregarded its mechanical role in the infected cell AFM simulations. Additionally, we postulated that most of the changes seen in Young’s modulus were due to the morphological and structural alterations of the nucleus rather than the cytoplasm. First, we examined the effects of changes in nuclear volume and height on the simulated AFM results while keeping everything else constant (Fig 5B, see also S2 Fig). While there was no change in the simulated Young’s moduli between the noninfected and 8 hpi similar-sized nuclei, the increase in the nuclear size at 12 hpi led to a significant increase in the simulated stiffness compared to the noninfected nuclei. Next, we simulated the effect of chromatin marginalization on the simulated Young’s moduli and found no significant change (Fig 5C). Finally, we investigated the importance of lamina-chromatin tethering by simulating the 8 hpi nuclei with and without LADs. We found that the omission of LADs significantly reduced Young’s modulus (Fig 5D). However, this decrease in stiffness was relatively minor compared to the experimentally observed change between noninfected and 8 hpi nuclei. Therefore, since these changes in the nucleus could not explain the experimentally observed reduction in nuclear stiffness, other mechanical changes must occur during HSV-1 infection.

To understand the changes in nuclear mechanics that could reduce nuclear stiffness, we focused on two key changes: decreased outward force and lamina disintegration (Fig 5E). The first change involves a decrease in cytoskeletal force on the nucleus via the LINC complexes or an osmotic pressure difference between the cytoplasm and the nucleus. The second change, lamina disintegration, a phenomenon observed during HSV-1 infection, was implemented by reducing the mechanical strength of a given percentage of connections in the lamina, without altering the number of LADs. Decrease in the outward forces had the most significant effect, as shown by the reduction of the simulated Young’s moduli (Fig 5F). Lamina disintegration was generally insignificant when the outward forces were high, but became more important at lower outward force strengths. Thus, our computational data suggested that outward forces, including cytoskeletal forces and osmotic pressure, had the most prominent effect on nuclear stiffness.

Next, we used the model to study whether the experimentally observed increase in NE membrane tension during the infection can co-occur with decreased outward forces and disintegration of the lamin network. We found that tension in the nuclear membrane is affected by both lamina disintegration and the decrease in outward forces (Fig 5G). Lamina disintegration increased nuclear membrane tension, whereas a reduction in outward forces decreased it. The increase in membrane tension during lamina disintegration can be explained by a larger share of the outward forces acting on the membrane rather than on the lamina.

Importantly, these results show that the combined effects of lamina disintegration and reduced outward forces can lead to a constant or increased NE membrane tension during infection. Furthermore, we compared the distribution of lamina tension in the NE as the lamina was disintegrated (Figs 5H and 5I). There were more local high-tension spots on the top surface of the nucleus as the lamina disintegrated, which led to an overall increase in the nuclear lamina tension. In all cases, the tension was higher on the sides of the nucleus (Fig 5H), which is explained by the absence of an actin cap to counteract the outward nuclear forces in these regions. Taken together, the simulation results indicate that the decrease in nuclear stiffness during HSV-1 infection is a consequence of reduced cytoskeletal forces and osmotic pressure, rather than of lamina disintegration. Also, reduced nuclear stiffness can coincide with increases in nuclear membrane and local lamina tension. Lastly, based on our assumptions and simulations, chromatin marginalization did not affect the nuclear stiffness during the infection.

### Intranuclear cytoskeletal pulling forces and cell-substrate contact area decrease in infection

Our computational modeling indicated that decreases in cytoskeletal pulling forces and osmotic pressure reduce nuclear softness. Previous studies have shown that suppressing the LINC complex protein SUN2 alters nuclear mechanical properties, leading to nuclear softening [23]. Consistent with our studies, the infection-induced decrease in SUN2 has been detected in HSV-1 and human cytomegalovirus infections [57,58]. Nuclear Ca^2+^ ions are connected to the shape regulation of the nucleus [59], nuclear translocation [60], and chromatin compaction [61]. Here, to monitor virus-induced nuclear changes and to verify the reliability of the model, we performed experimental validation of mechanical force on the LINC complex and of nuclear Ca^2+^ levels during infection.

First, we analyzed the effect of HSV-1 infection on the mechanical forces transmitted through the cytoskeleton to the nucleus. Here, we used a nesprin FRET tension sensor with a phasor approach to FLIM to measure the force on the LINC complex [62]. These studies indicated that the cytoskeletal pulling force of the nucleus was significantly lower in infected Vero cells at 8 and 12 hpi compared to noninfected cells (Figs. 6A and B, see also S5 Fig). This demonstrates that the mechanical linkage of the cytoskeleton to the nucleus is altered in infection, and the LINC complex forces decrease.

**Figure 6.**
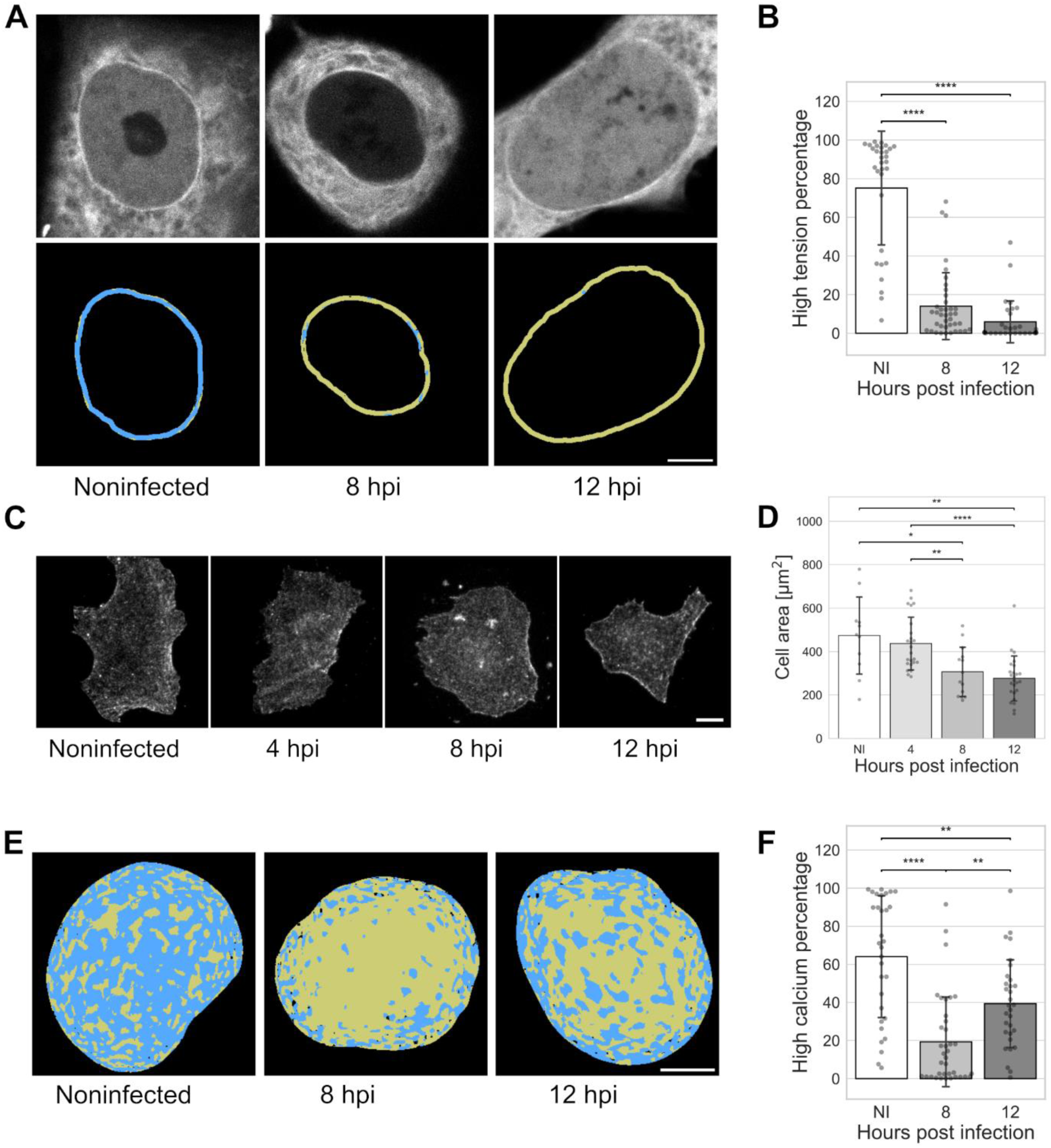
Cytoskeletal forces on the nucleus decrease in infection. **(A)** The nesprin tension FRET sensor analysis of the cytoskeletal pulling force in noninfected and infected Vero cells at 8 and 12 hpi. FRET donor fluorescence intensity (grey) and nesprin tension, quantified from the donor lifetime (S5 Fig), divided into low (yellow) and high (blue) tension regions at the segmented nuclear envelope. Scale bar, 5 μm. **(B)** The percentage of high tension regions in the segmented nuclear membrane in noninfected (NI) and infected cells at 8 and 12 hpi (n=31, 39, and 31, respectively). **(C)** Sum projection of beta-actin immunolabeled noninfected and infected cells at 4, 8, and 12 hpi. Scale bar, 10 μm. **(D)** The measurement of the area covered by noninfected and infected cells at 4, 8, and 12 hpi (n = 11, 24, 14, and 25, respectively). **(E)** Phasor FLIM analysis of calcium concentration using a single-molecule calcium biosensor in noninfected and infected Vero cells at 8 and 12 hpi. The calcium concentration in the nucleus divided into low (yellow) and high concentration (blue) regions according to each pixeĺs fluorescence lifetime distribution in the phasor plot (S6 Fig). Scale bar, 5 μm. **(F)** The percentage of high calcium concentration regions of the total nuclear region in noninfected cells and infected cells at 8 and 12 hpi (n=31, 37, and 31, respectively). The error bars show the standard deviation. Statistical significance was determined using Tukey’s test, and the significance values are denoted as **** (p<0.0001), ** (p<0.01), or * (p<0.05). Nonsignificant differences (p ≥ 0.05) are not labeled.

Next, we labeled cells for beta actin and measured their contact areas to the substrate in noninfected and infected cells. The analysis revealed that the cell contact area was reduced during infection (Figs 6C and D). This supports the idea that a decrease in outward forces due to reduced actin cytoskeleton pull occurs during the infection.

Finally, we measured nuclear Ca^2+^ concentration using phasor analysis for FLIM of single-molecule calcium biosensor G-Ca-FLITS [63,64]. The data show that nuclear Ca^2+^ concentration is lower in infected Vero cells at 8 and 12 hpi than in noninfected cells (Figs 6E and F, see also S6 Fig). Altogether, the reduced nuclear Ca^2+^ concentration suggests that chromatin compaction may decrease in infection, which probably affects nuclear mechanics.

## Discussion

Previous studies on isolated nuclei have shown that nuclear stiffness increases after the nuclear injection of the HSV-1 genome [65]. Our data shows that this is a transitory phenomenon or specific to isolated nuclei since the progression of the infection in living cells led to nuclear softening. The observed softening could be explained by changes in nuclear structure caused by the enlarged VRC at the center of the nucleus, changes in the nuclear lamina and chromatin, a decrease in the external pull from the cytoskeleton, or a decrease in osmotic pressure.

The progression of HSV-1 infection leads to an enlarged nuclear VRC, and at the late stages of infection, the VRC can occupy one-third of the nuclear volume [28]. Our SXT results indicate that the VRC region is less dense than the euchromatin region in noninfected cells and the marginalized chromatin in infected cells. This shows that HSV-1 infection forms an expanded low-density nuclear center surrounded by chromatin. It has been demonstrated that particle density is proportional to stiffness in many materials, such as collagen-based hydrogels [66,67]. While other factors, such as crosslink density between hydrogel fibers [68], also influence stiffness, the lower density of the VRC may indicate lower stiffness than chromatin and contribute to nuclear softening during infection. It has remained unclear how the infection and the enlarged VRC affect intranuclear ion and osmotic concentration. Modeling has stated that nuclear osmotic pressure is proportional to the macromolecular concentration [69,70]. Thus, the reduced macromolecular concentration in the VRC region likely contributes to regulating nuclear stiffness by decreasing osmotic pressure.

One of the major regulators of nuclear stiffness is the nuclear lamina associated with peripheral heterochromatin, particularly lamins A and C [10,12,13,71–74]. Studies with lamin A/C-deficient yeast cells have shown that chromatin tethering to the nuclear membrane increases the mechanical stiffness of the NE, and untethering of the chromatin leads to a softer nucleus [7]. In HSV-1 infection, nuclear softness may be altered by virus-induced modification of the nuclear lamina. Herpesvirus infection does not change the expression of lamins, but results in local dispersion of the nuclear lamina by phosphorylation and redistribution of lamins A/C and B. Various viral proteins and kinases (UL13, UL34, ICP34.5, and Us3) mediate these processes [39,75–78]. We showed how the mobility of the nuclear lamina decreases in infection, suggesting a possible increase in its tension or rigidity. However, it cannot be ruled out that infection-induced changes in the microtubules might also play a role in NE mobility [7]. Our simulations showed that the effect of lamina disintegration alone was not significant on simulated nuclear stiffness. Still, it became relevant when the outward forces, described as the combined effect of cytoskeletal pulling forces and osmotic pressure on the NE, were reduced. Finally, we showed that nuclear membrane tension increases at late infection, coinciding with nuclear volume expansion. This could be explained by earlier studies showing that nuclear swelling induces mechanical stretching forces that elevate membrane tension [79]. Membrane tension could also be affected by changes in lipid deformation during the formation of the viral egress complex [80]. Our simulations showed that the disintegration of the lamina led to increased membrane tension even when outward forces were reduced, because more of the outward forces were subjected to the membrane rather than to the lamina.

Our simulations were conducted to explain the experimental softening of the nucleus. They indicated that a reduction in outward forces led to the most significant decrease in nuclear stiffness, suggesting this could be the primary mechanism underlying the experimentally observed change in nuclear stiffness during the infection. The outward forces described the combined effects of cytoskeletal forces on the NE and osmotic pressure in the nucleus. Nuclear elasticity is partly regulated by the cytoskeletal tension mediated by the LINC complex, which links the nucleo- and cytoskeleton [10,74,12]. Our data showed that the force transmission from the cytoskeleton to the nucleus is altered in HSV-1 infection. First, we observed that SUN2 expression decreases during HSV-1 infection. Suppressing LINC protein SUN2 changes the mechanical properties of the nucleus and causes nuclear softening [23]. In macrophages, a deficiency of SUN1 and SUN2 proteins causes a decrease in nuclear size and an increase in nuclear elasticity [81]. SUN2 is a short-lived protein with a degradation half-life of 240 min [82], and temporal proteomics analysis has confirmed the reduced amount of SUN2 at late HSV-1 infection [57]. This most likely regulates the mechanical properties of the nucleus and reduces nuclear stiffness as the infection progresses. However, it cannot be the sole effector, since nuclear stiffness is already lower at 4 hpi, when SUN2 has not yet been downregulated. SUN2 depletion also modifies the NE double membrane and the perinuclear space [82,83]. Second, we showed that tension on nesprin proteins decreases in the infection. It is known that the force on the LINC complex can directly affect nuclear elasticity, and a decrease in the cytoskeletal pull softens the nucleus [22,23]. Additionally, it is known that HSV-1 rearranges the actin cytoskeleton via cofilin-1 [84]. Finally, we observed that the cell-substrate contact area decreased during infection, supporting the observed reduction in cytoskeletal tension [85]. Together, the downregulation of SUN2, the observed decrease in cell-substrate contact area, and the reduced force on the LINC complex indicate that the cytoskeletal pulling force on the nucleus was reduced.

The subdiffusive capsid motion in the nucleus is restricted by chromatin marginalized toward the NE. However, later in infection, the chromatin barrier becomes more permissive, and the probability of capsids entering the chromatin increases [33]. The mechanical changes that we observed might partly explain this phenomenon. Specifically, downregulation of the SUN protein has been shown to reduce chromatin condensation [86]. In HSV-1 infection, decreased SUN2 expression could facilitate the progression of infection and the nuclear egress of HSV-1 capsids, possibly by remodeling chromatin and NE organization [87]. The recent cytomegalovirus data show that changes in LINC-complex-mediated microtubule mechanotransduction lead to reorganization of both hetero- and euchromatin [88]. Moreover, nuclear chromatin compaction is also regulated by Ca^2+^ ion concentration. At low calcium levels, the negative charges of DNA remain partially unneutralized, leading to more expanded fibrous chromatin [61]. The lower calcium concentration that we observed during infection may therefore trigger local remodeling and decondensation of chromatin and possibly affect chromatin dynamics, which could explain the facilitated capsid diffusion through the chromatin during the nuclear egress of viral capsids. The decreasing calcium concentration also suggests that intranuclear osmotic pressure may decrease during infection. However, calcium is only one of the many ions and molecules that affect osmotic pressure, and we cannot make conclusions based solely on it. Moreover, due to the NLS-tagged sensor we used, we cannot assess the difference in calcium concentration between the nucleus and the cytoplasm.

In summary, herpesvirus infection and the emergence of enlarged, low-density VRCs lead to changes in nuclear volume, organization of chromatin and the nuclear lamina, and the connections to the cytoskeleton. Virus-induced nuclear deformation alters the nuclear environment and triggers changes in the intrinsic forces acting on the nucleus (Fig 7). Altogether, these profound changes cause nuclear softening. Our simulations show that the measured changes in the nuclear lamina and size have a relatively minor effect on stiffness compared to nucleocytoplasmic forces, namely cytoskeletal-nuclear connections and osmotic pressure. While the reorganization of the lamina is known to be essential for the nuclear egress of new HSV-1 viral capsids, the role of the predicted reduction in nucleocytoplasmic forces during infection remains elusive. Changes in the osmotic balance between the nucleus and the cytoplasm can, for example, originate from changes in nuclear transport, which HSV-1-induced changes in the nuclear pore function could cause [89]. These emerging insights shed new light on the virus-induced fundamental remodeling of nuclear organization and mechanobiology, and their impact on the progression of HSV-1 infection.

**Fig 7.**
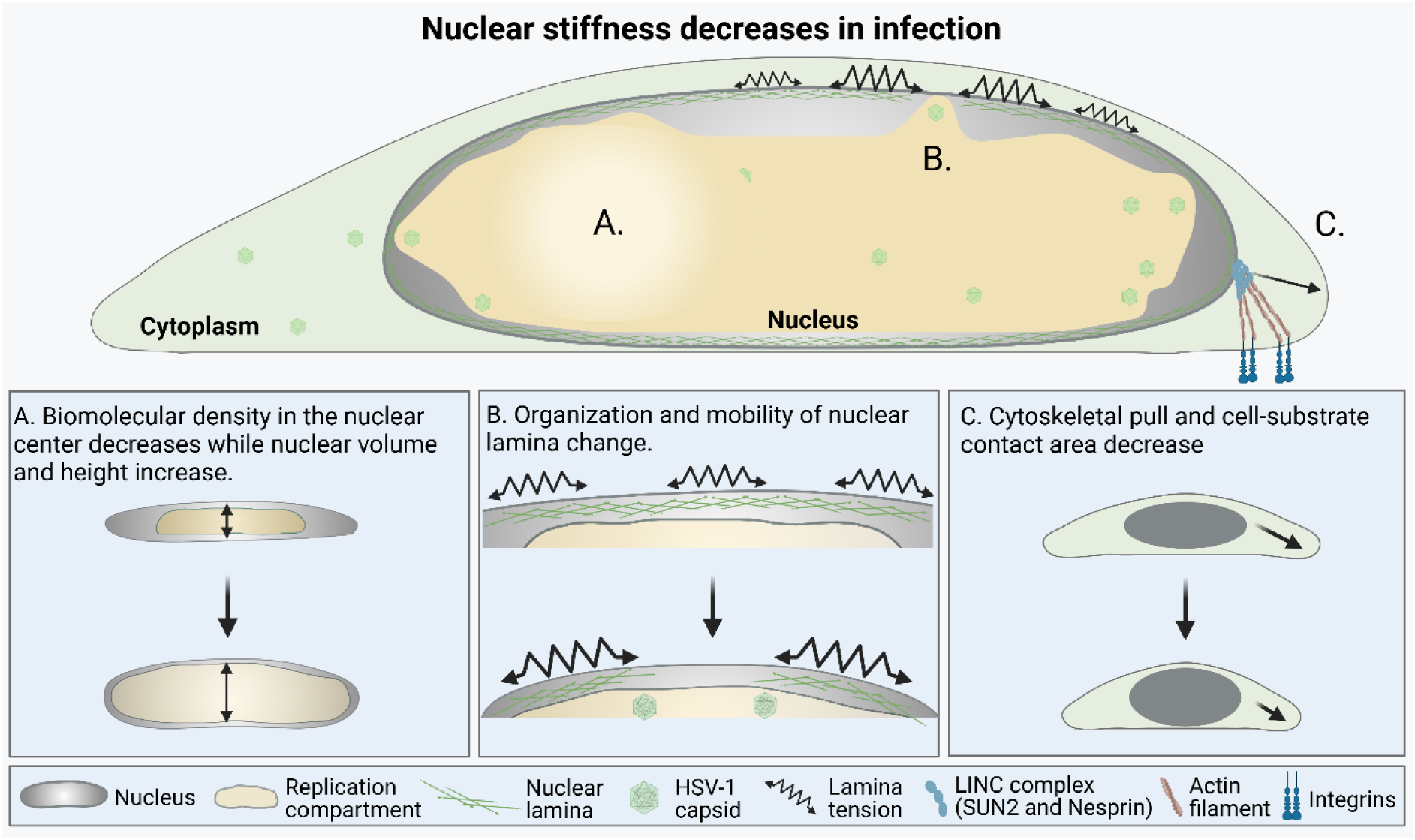
Virus-induced changes in mechanical forces regulating nuclear stiffness. HSV-1 infection leads to extensive changes in nuclear organization and mechanical forces. **(A)** The infection results in an enlarged nuclear viral replication compartment, a low-density area surrounded by a condensed chromatin layer. **(B)** Changes in the organization and mobility of the nuclear lamina accompany the increase in nuclear volume and height. **(C)** Nuclear stiffness is affected by cytoskeletal pull. The key regulator of cytoskeletal tension of the nucleus, LINC complex protein SUN2, is strongly downregulated in infection, and nesprin tension decreases together with the cell-substrate contact area. Viral infection alters the mechanical properties of the nucleus and leads to nuclear softening.

## Materials and Methods

### Cells and viruses

Mouse embryonic fibroblast cells (MEF, ATCC^®^ CRL-2991™) and African green monkey kidney cells (Vero, ATCC^®^ CCL-81™) were grown in Dulbecco’s modified Eagle medium (DMEM) supplemented with 10% fetal bovine serum, L-glutamine, and penicillin-streptomycin (Gibco*-*Invitrogen, Carlsbad, CA) at 37 °C in the presence of 5% CO_2_. HA-tagged progerin, a truncated form of lamin A [90], was expressed in Vero cells using adenovirus [91]. Lamin A adenovirus was used to overexpress wild-type lamin A in Veros. Lamin A adenovirus (based on RefSeq BC014507) was purchased from Vector Biolabs and used as previously described [91]. The cells were infected 4, 8, or 12 hours before imaging and measurements with the wild-type 17+ strain, EYFP-ICP4 (vEYFP-ICP4 [92]), or VP26-mCherry virus (HSV1(17^+^)Lox-CheVP26 [93–95]) at a multiplicity of infection of 5.

### Atomic force microscopy (AFM)

Force spectroscopy AFM experiments were performed on noninfected and infected Vero cells at 4, 8, and 10 hpi using a NanoWizard 4 XP instrument (JPK BioAFM, Bruker Nano GmbH, Berlin, Germany) mounted on an inverted microscope (Nikon Ti-U, Nikon Instruments Europe B.V., Amsterdam, the Netherlands) equipped with a standard monochrome CCD camera (ProgRes MFCool, Jenoptik, Jena, Germany). Fluorescence imaging was performed using wide-field illumination (Intensilight Hg lamp, Nikon) with a 100X objective (Nikon CFI Apo VC, 1.4 NA, oil immersion) and the appropriate filter cube for EYFP. A software module (DirectOverlay, JPK Bio-AFM, Bruker Nano GmbH, Berlin, Germany) calibrated the tip position with the optical image. Nanomechanical measurements were performed in spectroscopy mode using a 6.62 μm probe (CP-PNPL-SiO-C from SQube, mean cantilever spring constant k_cant_ = 0.06 N/m). Before each set of acquisitions, the sensitivity and spring constant of the cantilever were calibrated (thermal noise method). The measurements were performed in buffered culture medium at 37°C under 5% CO2, with a relative setpoint of 2 nN, a Z-length of 12 µm, and a constant extension velocity of 1 µm/s. The center of the nucleus of a cell was visualized by fluorescence, and ten successive force curve measurements were taken. Raw data were cleaned manually by removing any aberrant contact points, and each force curve was fitted with a Hertz model to calculate Young’s modulus using JPK data Processing software (7.0.181). The average pressing depth ranged from 0.9 μm (noninfected) to 3.3 μm (10 hpi), as analyzed by randomly selecting about 50% of cells at each infection time point. This reflects the decrease in nuclear stiffness that occurs during the infection. The average measures were plotted by cell (one point is one cell) after filtering out outliers (z-score>3).

### Cryo-soft X-ray tomography (SXT) of suspended cells in capillaries

MEF cells were seeded into culture flasks (Corning, Corning, NY) and infected with HSV-1 vEYFP-ICP4 at an MOI of 5 for 4 and 8 hours at 37**°**C. Noninfected and infected cells were detached with trypsin-EDTA, pelleted by centrifugation (300x*g*), and washed with PBS. Cells were then fixed with glutaraldehyde (0.5%) and paraformaldehyde for 10 minutes at RT and 35 minutes at 4°C (4%, cat. No. 15713-S; Electron Microscopy Sciences, Hatfield, PA), washed once with PBS, pelleted by centrifugation (300x*g*), re-suspended in the PBS buffer, filtered, loaded into thin-walled cylindrical borosilicate glass capillaries (Hilgenberg GmbH, Hilgenberg, GER, Cat. No. 4023088), and then vitrified by quickly plunging them into liquid-nitrogen-cooled (∼90 K) liquid propane. SXT tomographic data were collected by full-rotation imaging with a soft X-ray microscope (XM-2) in the National Center for X-ray Tomography at the Advanced Light Source (http://www.als.lbl.gov) of Lawrence Berkeley National Laboratory (Berkeley, CA). Cells were kept in a stream of liquid-nitrogen-cooled helium gas to avoid radiation damage during data collection. For each data set, 180° rotation tomographs (1 projection image per 2° angle) were collected [96,97]. Projection images were aligned by tracking fiducial markers and reconstructed using automatic reconstruction software [98]. The high- and low-density chromatin was segmented using histogram-based separation into two classes in Dragonfly (Comet Technologies Canada Inc., Montreal, Canada).

### Focused Ion Beam Scanning Electron Microscopy (FIB-SEM)

The volume imaging data set of MEF cells with enhanced chromatin contrast was generated using FIB-SEM. MEFs, noninfected or HSV-1 infected at an MOI of 5 for 8 h, were processed for imaging as described earlier [99]. The Chrom EM method was used to enhance the contrast of DNA [45]. The deposition of the osmiophilic polymer of diaminobenzidine, specifically on the DNA, was carried out using the fluorescent emission of DRAQ5. For this, the cells were placed on a cooled stage of a Leica DM IL LED microscope and exposed to 620 ± 30 nm light in the presence of diaminobenzidine, inducing its local polymerization over DRAQ5-bound DNA. The cells were treated with osmium tetroxide, followed by dehydration in a graded series of ethanol and embedding in resin. One set of cells was thin-sectioned and then poststained with 2% uranyl acetate and lead citrate. Another set of cells was coated with platinum. The sections were imaged using a Jeol JEM-1400HC transmission electron microscope with an Olympus SIS Quemesa bottom-mounted CCD camera (80 kV). The datasets were acquired in a Zeiss Crossbeam 500 using backscattered and secondary electron signals. The area of interest was exposed using a milling beam (0.7 nA at 30 kV) and imaged with a scanning electron beam (0.31 or 0.50 nA, 1.5 or 1.6 keV). The milling and scanning process of the cell cross-section produced an image volume with a voxel size of 2.5 x 2.5 x 5 nm^3^. The datasets were collected and aligned using Zeiss Atlas 5 software with a 3D tomography module. Microscopy Image Browser [100] was used to segment the capsids, chromatin, and the NE. The chromatin segmentation was done semi-automatically using MIB (https://mib.helsinki.fi/) and cleaned using iterative intensity-based erosion. The local thickness of the chromatin was measured in ImageJ (https://imagej.net/imagej-wiki-static/Local_Thickness) using Python; local maxima were identified, and the local thickness value was extracted. The distance of these local maxima to the NE was then calculated using the Euclidean distance transform.

### Confocal imaging and analysis

Wild-type Vero cells, Vero cells expressing HA-tagged progerin, or Vero cells overexpressing wild-type lamin A were grown on glass coverslips for microscopy to analyze nuclear volume. The cells were infected with HSV-1 EYFP-ICP4 and fixed at 8 and 12 hpi with 4% PFA, and DNA was labeled with DAPI. Viral EYFP-ICP4 was boosted with an anti-GFP antibody. Cells were embedded into VECTASHIELD® PLUS Antifade Mounting Medium (H-1900) (Vector Laboratories). The cells were imaged using a Leica SP8 confocal microscope with a UPLSAPO 63x oil immersion objective (NA 1.35). DAPI (Invitrogen) was excited with a 405 nm diode laser, and a 410-439 nm band-pass filter was used to detect fluorescence. EYFP was excited with a 488 nm laser, and the fluorescence was collected with a 519-540 nm bandpass filter. Lamin A/C was excited with a 653 nm laser, and fluorescence was collected with a 661-839 nm filter. Stacks of 1024 x 1024 pixels were collected with a 120 nm/pixel pixel size in the x- and y- directions and 400 nm in the z-direction. The nuclei were segmented using the minimum cross-entropy threshold algorithm [101] for the DAPI channel.

For analyses of the nuclear intensity of lamins, Vero cells infected with HSV-1 EYFP-ICP4 and fixed at 8 and 12 hpi were labeled with antibodies against lamin A/C (mouse monoclonal antibody MAb; ab8984, Abcam, Cambridge, UK) and Lamin B1 (rabbit polyclonal antibody, rAb; ab16048, Abcam). The primary antibodies were followed by goat anti-rabbit or anti-mouse Alexa 546- (lamin A/C) and 647- (lamin B1) conjugated secondary antibodies (Thermo Fisher Scientific, Waltham, MA, USA). DNA was labeled with Hoechst 1:10000 in PBS for 10 min. Cells were embedded into VECTASHIELD® PLUS Antifade Mounting Medium (H-1900) (Vector Laboratories). The immunolabeled cells were imaged using a Leica TCS SP8 FALCON laser scanning confocal microscope (Leica Microsystems, Mannheim, Germany). Alexa 546 and Alexa 647 were excited with 557 nm and 653 nm wavelengths of pulsed white light laser (80 MHz), respectively. The emission detection range was 568-648 nm for Alexa 546 and 663-776 nm for Alexa 647. DAPI was excited with a 405 nm diode laser, and emission between 415-495 nm was detected. Laser powers for all channels were fixed. Stacks of 1024 x 1024 pixels were collected with 120 nm/pixel in the x- and y- directions and 240 nm in the z-direction. The nuclei were segmented using a minimum cross-entropy threshold algorithm [101] for the DAPI channel. The total nuclear intensity was calculated for the pixels belonging to the segmented nucleus, and the intensity near the NE was calculated by using only pixels having an Euclidean distance of 1 μm or less from the NE.

The distribution and mobility of lamins were imaged in living noninfected and HSV-1 mCherry-infected (8 and 12 hpi) Vero cells transfected with lamin A/C*-*Chromobody-GFP2 (Proteintech, Manchester, UK) by electroporation using Neon Transfection System (Thermo Fischer Scientific) with 1150 V and two pulses with a width of 20 ms. Before imaging, the cells were labeled with 1 µM Hoechst 33342. During the imaging with Leica SP8 Falcon, using Leica HC PL APO CS2 1.2NA 63X water immersion objective, Hoechst was excited with 405 nm laser, 1% laser power and the emission collected between 412-475 nm (HyD), the lamin A/C-Chromobody-GFP2 was excited with a 488 nm wavelength of white light laser, 10% laser power and the emission was collected between 495-560 nm (HyD), and the mCherry channel was excited using 587 nm, 10% laser power and emission was collected between 594-800 nm (HyD). The image size was 296 x 296 pixels with a 100 nm pixel size in the x- and y-directions and the imaging speed was bidirectional 1000 Hz with two sequences (1. Hoechst, 2. GFP and mCherry), resulting in 3.2 frames per second for 60 seconds. The lamina was analyzed by first removing its translational and rotational motion. The lamina was skeletonized at each time point, and the maximumintensity projection of the skeletons was taken. The area of the maximum intensity projection was calculated and divided by its length. The curvature was defined by discretizing the nuclear lamina using 16-pixel (1.6 μm) long line segments and calculating the angles between consecutive segments.

To measure the contact area of the cells during the infection, ICP4-EYFP-infected cells were labeled using a mouse Anti-beta Actin antibody ([mAbcam 8226], Abcam). The contact area of the cells during the infection was analyzed by taking the sum projection of the actin channel along the z-direction, thresholding the sum projection using the minimum cross-entropy algorithm [101], and calculating the area of the segmented region.

### Fluorescence lifetime imaging and analysis

Nuclear membrane tension was analyzed by FLIM using fluorescent membrane tension probe ER Flipper-TR^®^ (Spirochrome) in noninfected and HSV-1 17+ wt infected live Vero cells at 8 and 12 hpi. Measurements were performed with a Leica SP8 Falcon using Leica HyD SMD detectors and a Leica HC PL APO CS2 1.2NA 63X water immersion objective. The ER Flipper probe was used at 1 µM, and the cells were also labeled with Hoechst 33342 at 1 µM. One plane at the center of the nucleus, where the nuclear edges were clearly visible, was first acquired with 405 nm excitation and 415-465 nm emission to visualize DNA with Hoechst. Immediately after, the ER Flipper was imaged using 477 nm excitation and 550-650 nm emission, and line accumulation of 16. The scanning speed was 200 Hz, the repetition rate was 80 MHz, and the image size was 328x328 pixels with 81x81nm/pixel for both Hoechst and ER Flipper imaging. For tension analysis, pixel binning of 3 was used, resulting in approximately 500-1500 photons per pixel. After binning, a pixel-wise two-component fitting was performed, and the longer component, corresponding to membrane tension, was extracted. Using the Hoechst data, a 486 nm wide ring just outside the cellular DNA was used as a mask to obtain the nuclear membrane area from the tension data. The mean of the longer ER Flipper lifetime along the NE was calculated for each cell.

Pcdna nesprin TS, a gift from Daniel Conway (Addgene plasmid #68127; http://n2t.net/addgene:68127; RRID: Addgene #68127), was used to measure the amount of cytoskeleton tension towards the nucleus, reported by FRET, across the nuclear envelope. The Vero cells were transiently transfected by electroporation using the Neon Transfection System (Thermo Fischer Scientific) with two 1150 V pulses of 20 ms width, and grown overnight on a 35 mm live-imaging glass-bottom dish (Ibidi) in an antibiotic-free medium. The next day, or the day after, the cells were mock-infected and infected with HSV-1 VP26-mCherry, and imaged at 8hpi and 12hpi using a Leica SP8 Falcon at 37°C and 5% CO_2_, with a Leica HC PL APO CS2 1.2NA 63X water immersion objective. Before starting the imaging, the medium was changed to one without phenol red and containing 1 µM Hoechst 33342. All imaging was conducted approximately between 30 minutes before and 30 minutes after the designated time points, and the mock-infected cells were imaged in between the two time points. Mock-infected and infected cells were made from the same set of transfected cells for two biological replicates, and a third biological replicate of mock-infected cells was imaged for more n. From each imaged cell, first a reference image was taken from Hoechst (excitation 405nm, 1% laser power, emission 415-473nm, PMT), nesprin TS donor (excitation 470nm, 3% laser power, emission 485-500nm, HyD photon counting), nesprin TS acceptor (excitation 514nm, 3% laser power, emission 520-580nm, HyD photon counting) and mCherry (excitation 580nm, 3% laser power, emission 600-700nm, HyD photon counting) in four sequences, after which the FLIM-FRET data was acquired for the nesprin TS donor channel for 1 minute using frame repetition at the speed of 700 Hz and 80MHz pulse rate. Only cells with >50 photons per pixel at the NE were included in the analysis, as they provided a precise phasor cloud distribution without significant background contribution using the LAS X wavelet filter. On the other hand, cells with >1500 photons in the brightest pixel were excluded as the data was dominated by the high FRET of overexpressed non-bound nesprin TS. A pixel size of 69 nm (x,y), an image size of 360 (x,y) pixels, and a pinhole of 1 airy unit were used. An ROI for the NE was generated by automatic thresholding of nesprin TS FLIM or intensity data, or by manually drawing the NE in ImageJ. The data were divided into low- and high-FRET pixels at the NE, which inversely reports for high and low tension, respectively. A FRET efficiency threshold of 10% was used for the division. The number of pixels with low and high FRET at the NE was quantified for each cell, inversely representing the tension between the nucleus and cytoskeleton. A description of the phasor FLIM-FRET analysis workflow is presented in the S5 Fig.

To measure intranuclear calcium concentration, we used 3xnls-G-Ca-FLITS, a gift from Dorus Gadella (Addgene plasmid #191463; http://n2t.net/addgene:191463; RRID: Addgene #191463). The Vero cells were transiently transfected by electroporation using the Neon Transfection System (Thermo Fischer Scientific) with two 1150 V pulses of 20 ms width, and grown overnight on a 35mm live-imaging glass-bottom dish (Ibidi) in an antibiotic-free medium. The next day, or the day after, the cells were mock-infected and infected with HSV-1 VP26-mCherry, and imaged at 8hpi and 12hpi using Leica SP8 Falcon at 37°C and 5% CO_2_ and Leica HC PL APO CS2 1.2NA 63X water immersion objective. Before starting the imaging, the medium was changed to one without phenol red and containing 1 µM Hoechst 33342. All imaging was conducted approximately between 30 minutes before and 30 minutes after the designated time points, and the mock-infected cells were imaged in between the two time points. Mock-infected and infected cells were generated from the same set of transfected cells, with two biological replicates. The FLIM data were acquired by exciting the G-Ca-FLITS biosensor with a 474 nm laser at 3% power and 40 MHz repetition rate, for 1 minute at the speed of 1800 Hz. For increased photon detection rate, the emission spectrum was divided into two HyDs, first detecting 490nm-534nm and second 540-580nm, resulting in >400 photons per pixel. The 5 nm gap in the detection spectrum was because a PMT detector is placed in between the two HyDs in the system. Simultaneously to acquiring the G-Ca-FLITS FLIM data, Hoechst 33342 (excitation 405, laser power 0.5%, emission 415-490nm, PMT) and mCherry (excitation 586 nm, laser power 3%, emission 600-700nm, HyD) were acquired in separate sequences. A pixel size of 68 nm (x,y), an image size of 306 pixels (x,y), and a 1 Airy unit pinhole size were used. First, a control experiment was conducted with transfected noninfected cells, in which, during live FLIM acquisition, the medium was supplemented with 5X ionomycin (From APExBIO kit 2061). From this experiment, a calibration phasor FLIM line spanning low-to-high calcium concentrations could be generated, confirming the expected sensor function. Next, the phasor analysis of the biosensor FLIM data of noninfected and infected cells showed a clear lifetime difference along the ionomycin calibration experiment. Phasor cursors were drawn for these lifetime species to quantify nuclear pixels into high- or low-calcium concentration groups according to their fluorescence lifetime. The ratio of high and low-calcium concentration pixels for each nucleus was quantified. The phasor analysis of the G-Ca-FLITS FLIM data is further described in S6 Fig.

### Global Run-On and sequencing (GRO-seq)

GRO-seq analysis was performed as described previously [99]. Briefly, to prepare nuclei, noninfected and infected Vero cells (at 4 and 8 hpi) were swollen, and the nuclei were collected and stored at -80°C. The preparation of GRO-Seq libraries followed the previously described protocol [99,102]. The library quantification was performed using a Qubit fluorometer (Thermo Fisher Scientific) and prepared for 50 bp single-end sequencing on an Illumina HiSeq 2000 platform (GeneCore, EMBL Heidelberg, Germany). The raw sequencing reads were assessed by the FastQC tool (http://www.bioinformatics.babraham.ac.uk/projects/fastqc/) and the FastX toolkit (http://hannonlab.cshl.edu/fastx_toolkit/). GRO-seq reads were aligned to the ChlSab1.1 genome using the bowtie-0.12.7 software [103]. The best alignment was obtained with up to two mismatches and up to three locations per read. Differentially expressed genes were detected using ChlSab1.1.95.gtf coordinates, and the RNA.pl workflow of HOMER 4.3 and edgeR v3.2.2 were used for analysis. Genes were considered differentially expressed using the following cutoffs: log2 fold change 0.585, RPKM 0.5 in at least one sample, and adjusted p-value of 0.05. GRO-seq reads were mapped to the Human Protein Atlas (https://www.proteinatlas.org/) to identify lamins and nuclear lamina-related proteins. The interaction of these proteins was further mapped using the String database (https://string- db.org/) by selecting experimentally shown interactions using a threshold of 500. The non-interacting proteins were identified in the same String database, provided they interact with at least one detected protein. The resulting graph was generated in Python using the networkx package and then manually adjusted.

### Western blot

0.7 x 10^6^ Vero cells were plated on 5 cm cell culture dishes, grown over night, and infected the next day with HSV-1 17+ wt (MOI 5). Noninfected and infected cells (8 hpi and 12 hpi) weredetached by scraping. Scraped cells were collected in 1 ml of cold PBS and centrifuged at 2,000 x g for 5 min to pellet the cells. Cell pellets were resuspended in Laemmli sample buffer containing 2.5% 2-mercaptoethanol (Bio-Rad Laboratories) in PBS and heated at 95 °C for 10 minutes. Protein standards (Bio-Rad Laboratories) and 15 µl of samples were loaded onto 4– 20% Mini Protean TGX Stain-Free gels (Bio-Rad Laboratories), and run at 100 V for 80 minutes. Total protein stain in the gels was activated for 5 min, and the gels were imaged to detect the protein standards. Proteins were blotted onto 0.2 µm low fluorescence PVDF membranes with the Trans-Blot Turbo RTA Transfer Kit (Bio-Rad Laboratories) and Trans-Blot Turbo Transfer-System (Bio-Rad Laboratories). Blots and gels were imaged to verify protein transfer. After acquiring the total protein stain for each blot the membranes were blocked for 5 minutes in EveryBlot Blocking Buffer (Bio-Rad Laboratories, USA) and probed with primary antibodies against SUN2 (ab124916, Abcam), Lamin A (ab26300, AbcamPredicted band size: 74 kDa), Lamin B (ab16048, Abcam), and HSV-1 ICP4 (ab6514, Abcam). Primary antibodies were detected using fluorophore-conjugated secondary antibodies goat anti-mouse IgG StarBright Blue 520 and goat anti-rabbit IgG StarBright Blue 700 (Bio-Rad Laboratories). Gels and immunoblots were imaged with the Bio-Rad ChemiDoc MP Imaging System. Total band intensities for predicted-sized bands were quantified using Image Lab 6.1 software (Bio-Rad Laboratories).

### Computational modeling

The developed computational model described the nucleus as an NE, including the nuclear lamina, and chromatin components in three dimensions. The main model assumptions were: (1) the mechanical properties of the nucleus are dictated by the nuclear envelope, chromatin, and outward forces, including actin cytoskeleton prestress and osmotic pressure difference; (2) no changes in the mechanical properties occur during the short AFM measurement; and (3) the VRC has low density and has no effect on the AFM measurement. The NE was triangulated into a viscoelastic shell, and self-avoiding polymer chains were used to model the chromatin. Furthermore, additional interactions were defined to describe various connections, such as LADs between chromatin and the lamina, and between different parts of the chromatin. The herpesvirus VRC was described as a single coherent entity within the nucleus. It was represented similarly to the NE as a triangulated shape that repulses the chromatin as it grows.

The model was simulated by calculating a force affecting each vertex originating from various, usually Hookean, interactions and moving them using the damped equation of motion:

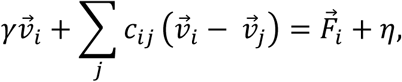

where *γ* is the system viscosity, *v*_*i*_ is the velocity of vertex *i*, *c*_*ij*_ is the friction between vertices *i* and *j*, *F*_*i*_ is the total force affecting vertex *i*, and *η* is the random thermal motion. The equation was solved using the conjugate gradient method, and the vertex movements were computed using the forward Euler method.

To simulate the AFM experiments, a cantilever with properties mirroring those of the one used in the experiments was used to push down on the nucleus, and the resulting force and indentation depth data were used to calculate the Young’s modulus using the Hertz model. The computational model was implemented using Julia programming language (v1.10.2), and the model and the parameters are described in more detail in S1 Text. The code is available on GitHub (https://github.com/atervn/nuclear_mech) and Zenodo (https://zenodo.org/records/14512358). The data fitting for the Hertz model was done using Matlab (The MathWorks Inc, R2021a, Natick, Massachusetts) and 3D visualization using ParaView [104].

## Data availability

The authors declare that the GRO-Seq data were deposited into the Gene Expression Omnibus database under accession number GSE243613 and are available at https://www.ncbi.nlm.nih.gov/geo/query/acc.cgi?acc=GSE243613. All other relevant data are in the manuscript and its supporting information files.

## Acknowledgments

We thank the staff of the National Center for X-ray Tomography (NCXT) and Advanced Light Source (Lawrence Berkeley National Laboratory, Berkeley, CA) for providing cryo-SXT experiments. We also thank Helena Vihinen from the Electron Microscopy Unit (University of Helsinki, Institute of Biotechnology, Helsinki Institute of Life Science, Helsinki, Finland) for preparing and imaging FIB-SEM samples and Elina Mäntylä for preparing the GRO-Seq samples.

---

**S1 Movie. AFM analysis of nuclear stiffness.** Animation of AFM imaging of infected Vero cells at 4 hpi (left). Retraction curves of four measurements showing the force (deflection) versus cell height in living cells, retraction (cyan) and extension (blue) curves, and contact points (arrows) are shown (right). The vertical red line shows the progression of the measurement.

**S2 Movie. Infection-induced changes of the nucleus.** Animation of cryo-soft X-ray (SXT) tomography imaging data of noninfected (NI) and infected MEF cells at 8 hpi. The high- (blue) and low-density chromatin (yellow), as well as the viral replication compartment (VRC, green), are shown.

**S3 Movie. Chromatin organization in the nuclear periphery is altered in infection.** Animation of focused ion-beam scanning electron microscopy (FIB-SEM) imaging data of noninfected and infected MEF cells at 8 hpi. The color bar indicates high and low chromatin thickness (yellow-blue). The nuclear envelope (cyan) and viral capsids (grey) are shown.

**S4 Movie. Remodeling of nuclear organization during the growth of the viral replication compartment.** Expansion of VRC as the infection proceeds. The chromatin is shown in blue, VRC in green, lamina-associated domains (LADs) of the chromatin and chromatin crosslinks in cyan, and the NE in grey.

**S5 Movie. Simulated AFM measurement of the infected cell.** AFM measurement of the nuclear stiffness of an infected cell. The effect of the colloidal probe of the cantilever (white) over the nucleus is presented. The cell membrane and cytoplasm are not presented. The chromatin is shown in blue, the lamina-associated domains (LADs) of the chromatin and chromatin crosslinks in green, and the NE in grey. The VRC located in the central region of the nucleus is not shown.

**S1 Table. Transcription of nuclear lamins and lamin-associated proteins is regulated in response to viral infection.** The table presents the results of GRO-seq analysis showing upregulated (green-blue) or downregulated (red-orange) lamins and lamin-associated protein genes in infected Vero cells at 4 and 8 hpi.

**S1 Text. Description of the model and the parameters**. A detailed description of the model assumptions, equations, solution method, atomic force microscopy simulations and analysis, the fitting process, and the parameter values.

**S1 Fig.**
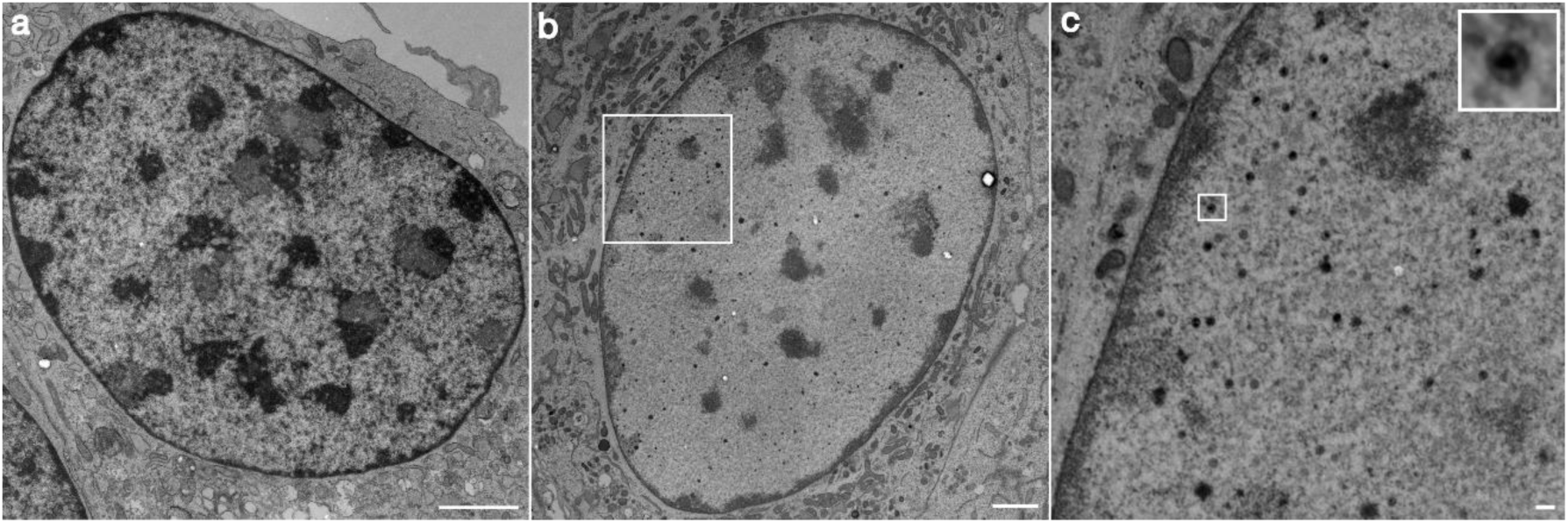
Distribution of nuclear chromatin and viral DNA in full capsids. Representative transmission electron microscopy images of **(A)** a noninfected and **(B)** an infected MEF cell at 8 hpi. ChromEM labeling was used to enhance DNA contrast. **(C)** Full viral capsids with DNA staining are shown. White squares show the magnified nucleoplasm and a full capsid. Scale bars, 2 µm (A, B) and 0.2 µm (C).

**S2 Fig.**
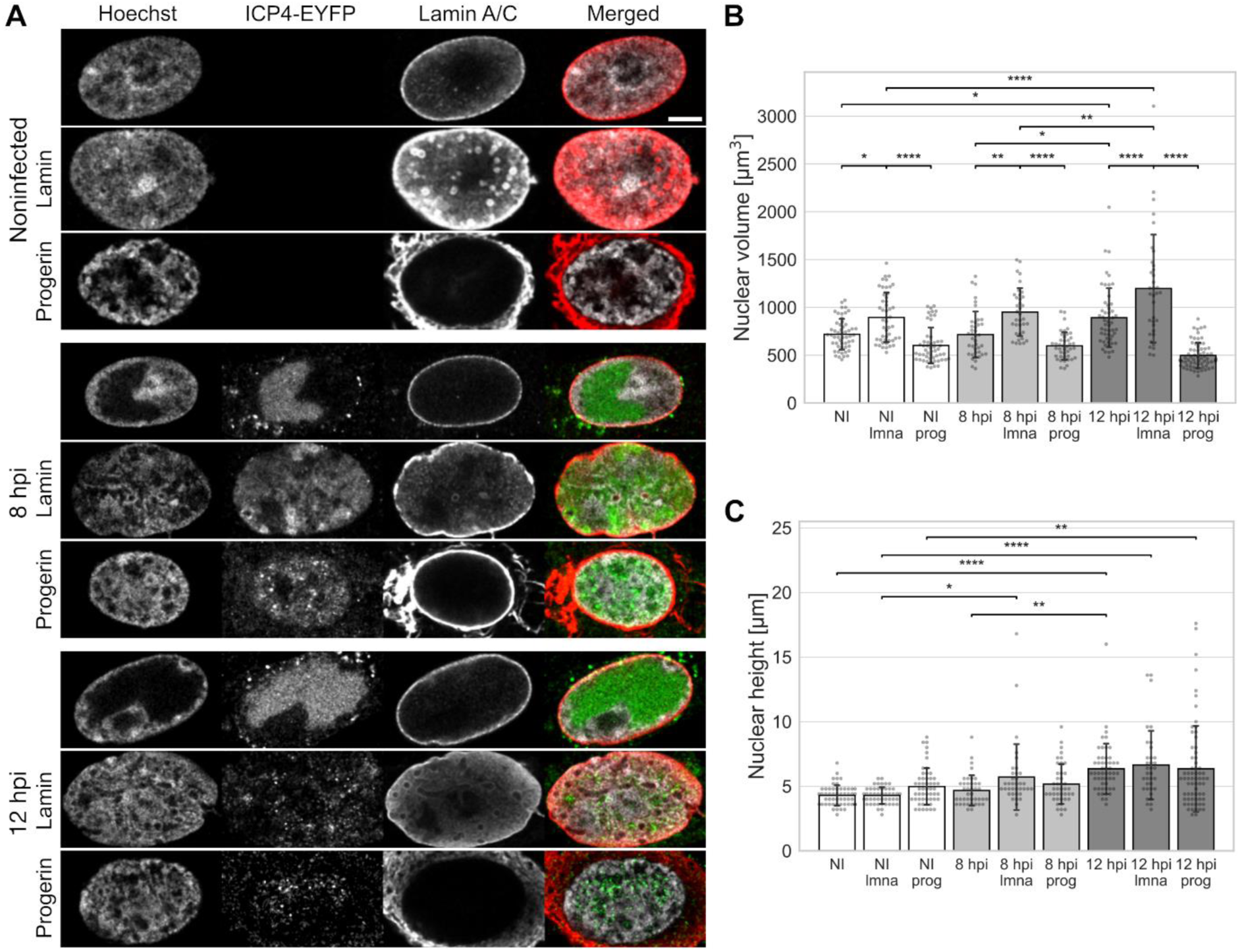
Overexpression of lamin A enhances nuclear growth. **(A)** Representative confocal images of noninfected and HSV-1 EYFP-ICP4-infected Vero cells at 8 and 12 hpi together with Vero cells overexpressing lamin A (lmna) and expressing progerin (prog). Lamin A/C was labeled with lamin A/C antibody (red), EYFP-ICP4 (green) shows the localization of the VRC, and the chromatin was labeled with Hoechst 33342 (grey). Quantitative analysis of **(B)** the volume and **(C)** the height of the nucleus in noninfected (NI) and infected cells (n=53, 48, 53, 40, 36, 42, 52, 35, and 67 for the bars from left to right). The error bars show the standard deviation. Statistical significance was determined using Tukey’s test, and the significance values are denoted as **** (p < 0.0001), ** (p < 0.01), or * (p < 0.05). Only values of the same time point or treatment were statistically compared. Nonsignificant differences (p ≥ 0.05) are not labeled.

**S3 Fig.**
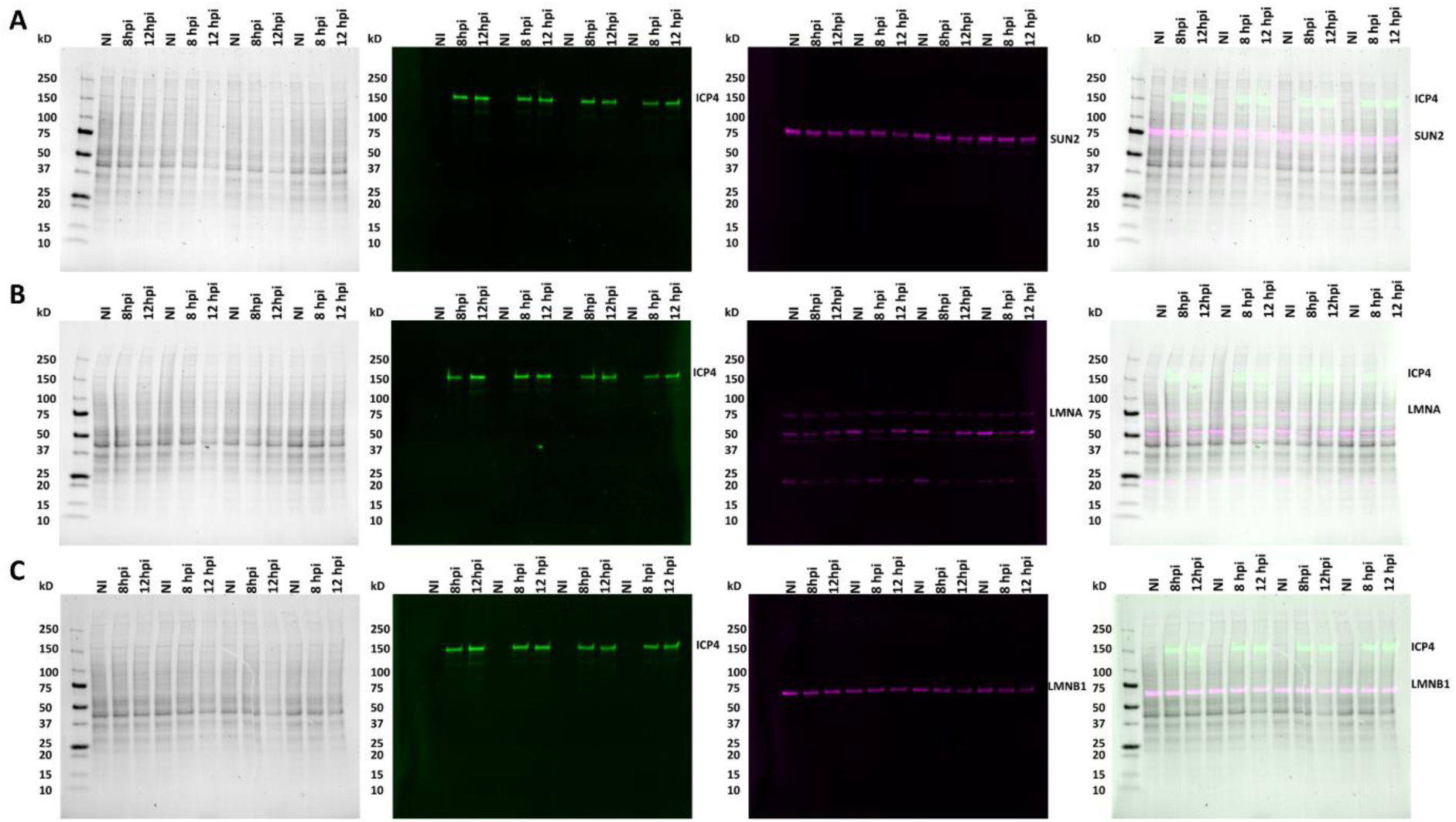
Expression of SUN2, lamin A, and lamin. **B**. Western blot of noninfected and infected (8 and 12 hpi) Vero cell lysates were analyzed for total protein and SUN2 **(A)**, LMNA **(B)**, or LMNB1 **(C)**. An antibody against ICP4 was used as a marker for viral infection. Each set of noninfected and infected lysates represents a replicate experiment (n = 4).

**S4 Fig.**
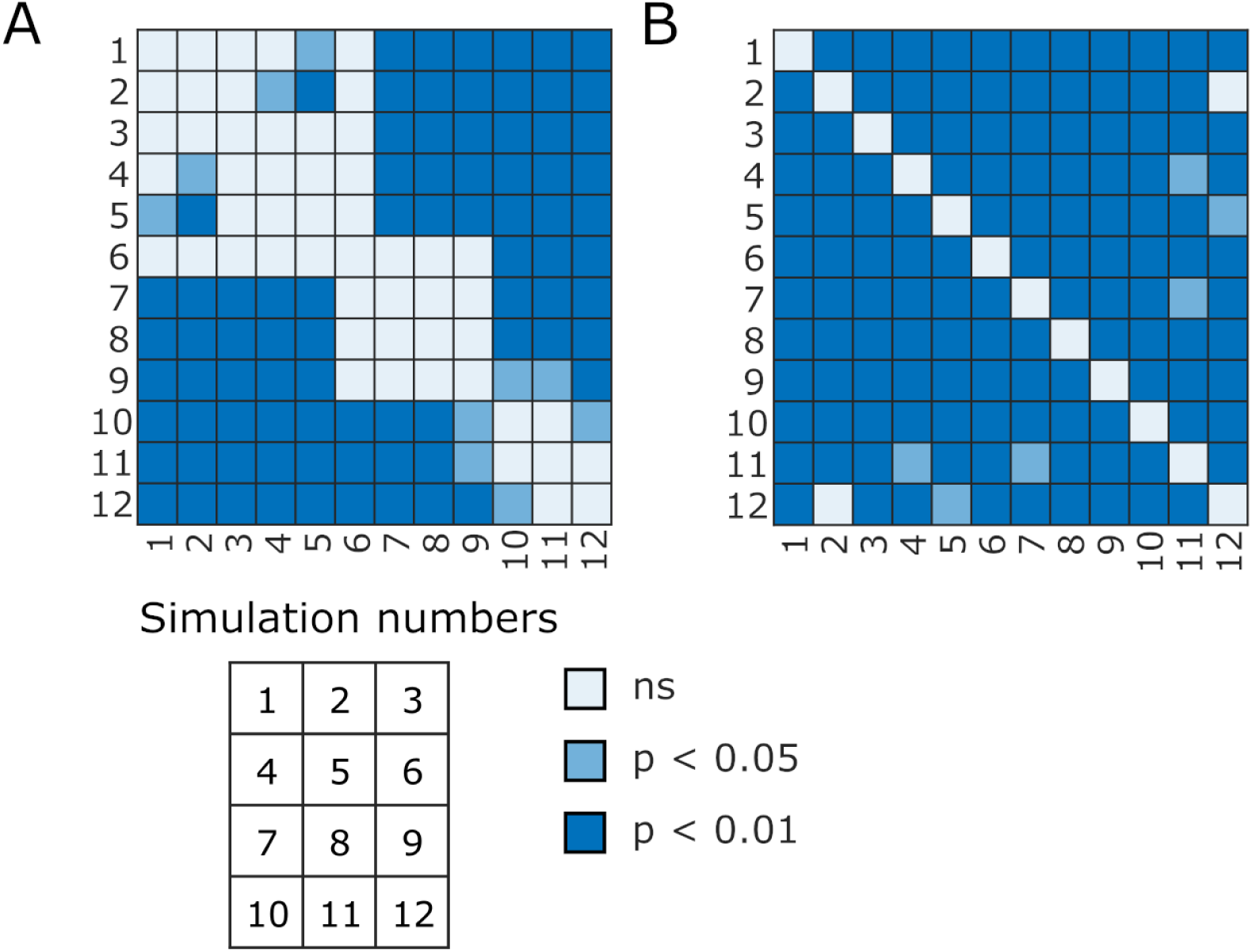
Statistical significances between simulations. Simulation case numbers and statistical significances between simulations presented in Figure 6 for **(A)** Young’s moduli (6F) and **(B)** nuclear envelope membrane tension (6G). Statistical significances were determined using the Games-Howell test. The significance values are denoted as p<0.01, p<0.05, or ns (not significant).

**S5 Fig.**
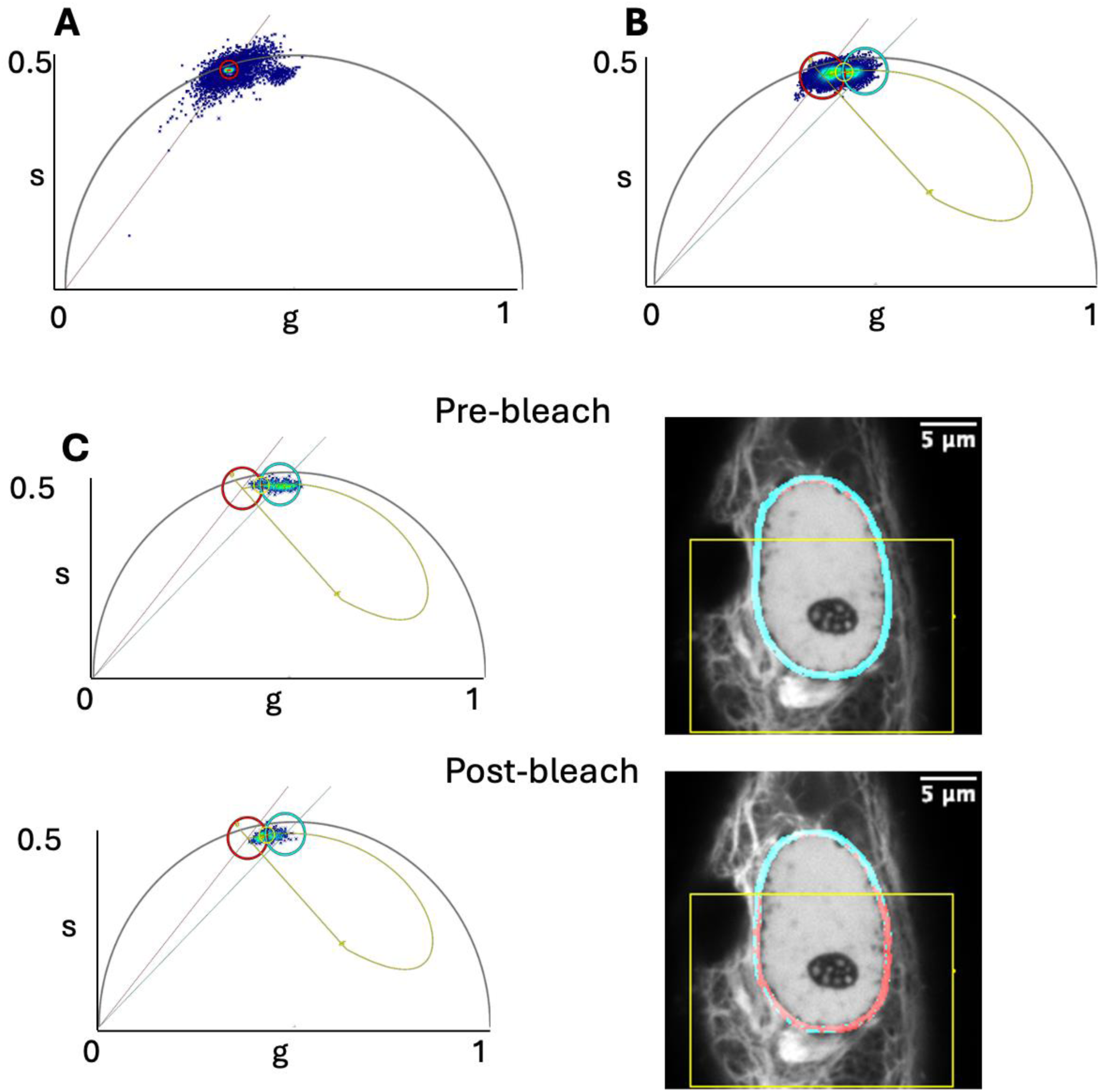
Tension sensor analysis of the mechanical forces transmitted to the nucleus. (**A**) Phasor plot analysis of the Nesprin TS donor fluorophore mTFP1 measured in noninfected and infected cells shows a lifetime distribution from 2.65ns to 2.71ns. (**B**) Phasor distribution of mTFP1 from FLIM measurement of noninfected and infected cells. The FRET trajectory was generated using LAS X (4.5.0) phasor analysis tool by setting the donor-only lifetime to 2.71 ns based on mTFP1 measurements, by locating the autofluorescence from non-transfected cells, and by adjusting the background contribution to ensure the trajectory passes through the measured phasor clouds. The division between low FRET/high tension (red phasor cursor at 2.47ns) and high FRET/low tension (cyan phasor cursor at 2.01ns) was set to 10% FRET efficiency (yellow phasor cursor). The pixels at the nuclear envelope (NE) of the measured cells were divided into low or high FRET areas according to their lifetime distribution in the phasor plot, and their relative amount was quantified. (**C**) An acceptor bleaching experiment to validate the Nesprin TS working in our system. FRET at the NE decreases after bleaching the acceptor from the area shown by the yellow box in a living cell.

**S6 Fig.**
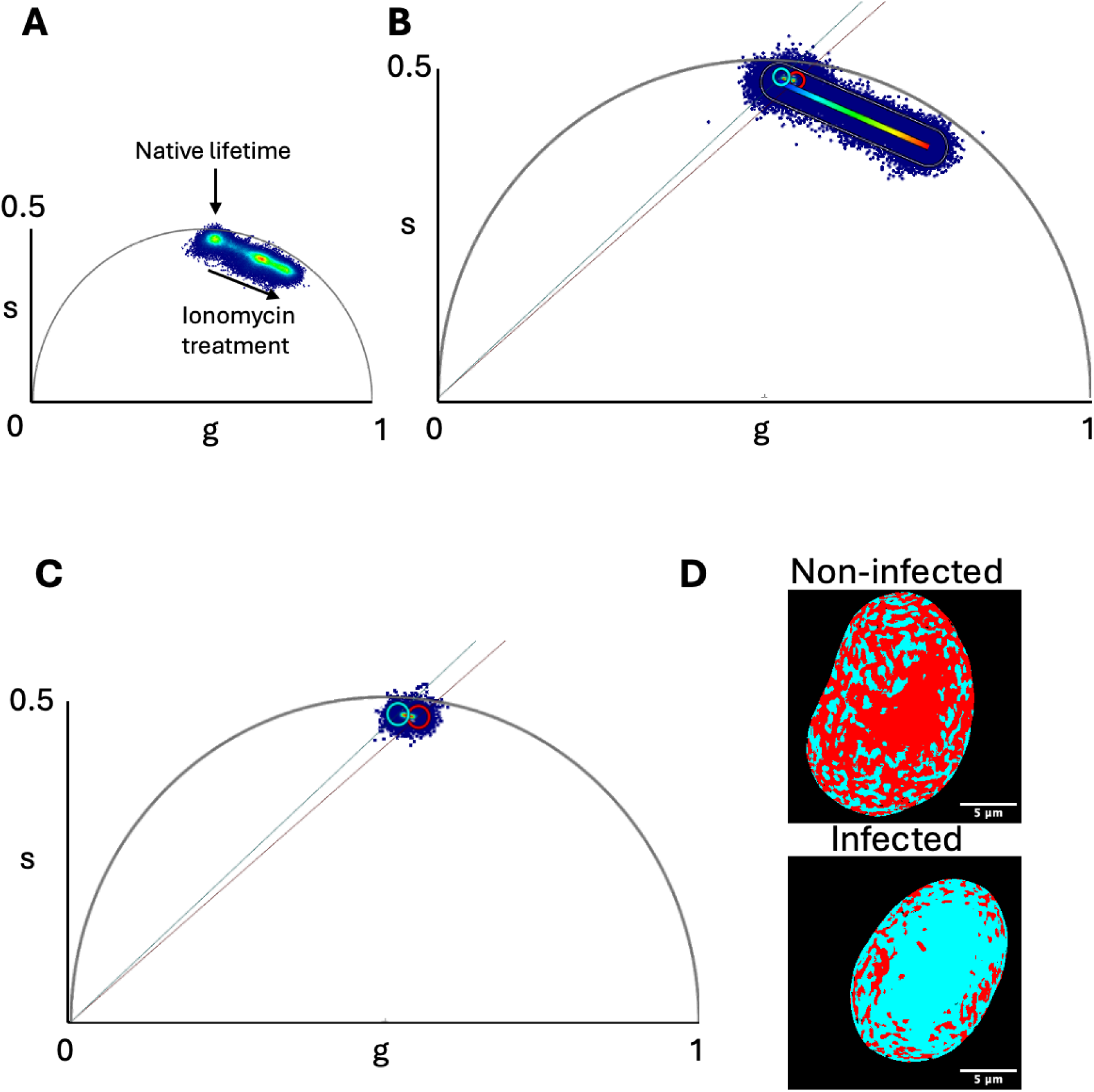
Detection of nuclear Ca^2+^ concentration using G-Ca-FLITS biosensor. **(A)** Phasor plot of the phasor distribution of live FLIM imaging of 3xnls-G-Ca-FLITS transfected cells. The phasor cloud in native cell state starts approximately from s=0.5, g=0.5 (top-center) and moves diagonally towards down-right during ionomycin treatment, confirming the expected shorter lifetime in high calcium concentration. **(B)** The calibration line (rainbow), generated according to the ionomycin measurement, demonstrated longer lifetime (cyan cursor) in infected cells than in noninfected cells (red cursor). This indicated reduced calcium concentration in infected cells. (**C**) Phasor distribution of noninfected and infected nuclei lifetime, and their division between low calcium (longer lifetime cyan phasor cursor set to 3.71 ns) and high calcium (shorter lifetime red phasor cursor set to 3.46 ns). (**D**) The pseudo-colored spatial distribution of pixels with high and low calcium concentration according to their lifetime distribution in the phasor plot in representative non-infected and 8 hpi nuclei.

